# Age-related susceptibility to grey matter demyelination and neurodegeneration is associated with meningeal neutrophil accumulation in an animal model of Multiple Sclerosis

**DOI:** 10.1101/2021.12.23.474008

**Authors:** Michelle Zuo, Naomi Fettig, Louis-Philippe Bernier, Elisabeth Pössnecker, Shoshana Spring, Annie Pu, Xianjie I. Ma, Dennis S. W. Lee, Lesley Ward, Anshu Sharma, Jens Kuhle, John G. Sled, Anne-Katrin Pröbstel, Brian MacVicar, Lisa Osborne, Jennifer L. Gommerman, Valeria Ramaglia

## Abstract

People living with multiple sclerosis (MS) experience episodic central nervous system (CNS) white matter lesions instigated by autoreactive T cells. With age, MS patients show evidence of grey matter demyelination and experience devastating non-remitting symptomology. What drives progression is unclear and has been hampered by the lack of suitable animal models. Here we show that passive experimental autoimmune encephalomyelitis (EAE) induced by an adoptive transfer of young Th17 cells induces a non-remitting clinical phenotype that is associated with persistent leptomeningeal inflammation and cortical pathology in old, but not young SJL/J mice. While the quantity and quality of T cells did not differ in the brains of old *vs* young EAE mice, an increase in neutrophils and a decrease in B cells was observed in the brains of old mice. Neutrophils were also found in the leptomeninges of a subset of progressive MS patient brains that showed evidence of leptomeningeal inflammation and subpial cortical demyelination. Taken together, our data show that while Th17 cells initiate CNS inflammation, subsequent clinical symptoms and grey matter pathology are dictated by age and associated with other immune cells such as neutrophils.

## Introduction

Multiple sclerosis (MS) is an autoimmune disease that causes demyelination of the central nervous system (CNS). MS is typically diagnosed in early adulthood as relapsing-remitting MS (RRMS)^1^. Disease activity waxes and wanes, with relapses characterized by lesion formation in the deep white matter initiated by infiltration of autoreactive T lymphocytes across the blood brain barrier^2^. Approximately 15-20 years following onset, or when symptoms are first manifested in people older than 40 years, the disease enters a progressive phase, becoming non-remitting^3^. Age is a major risk factor for disease progression in MS^4^ with epidemiological studies showing that older chronological age at onset is associated with faster time to disability milestones^3,5–7^.

A key hallmark of brain pathology in progressive MS is demyelination and neurodegeneration of the grey matter. Although present from the earliest stages of MS^8,9^, grey matter injury accrues with disease progression^10^, associates with motor deficits and cognitive impairments^11,12^, and extensive cortical damage at onset predisposes to a rapid transition into the progressive phase of the disease^13^. Unfortunately, the process of disease progression is ill-understood. Notably, immunomodulatory therapies that are effective at suppressing relapsing disease, have at best little impact on progression^14^.

It has been proposed that immunomodulatory therapies fail in progressive MS (PMS) because disease progression is not governed by immune cells^15^ despite immune cells being found in brain-adjacent regions of the progressive MS brain, in particular within the leptomeninges^16–18^. Moreover, these immune cells appear to be proximal to areas of grey matter injury^19–21^ and patients with aggregates of leptomeningeal immune cells harbour a number of pro-inflammatory molecules in their cerebrospinal fluid (CSF)^22^. Thus, it is likely that immune cells are involved in the clinical and pathological presentation of PMS, but therapies that are used in RRMS either do not target these immune cells, or there are redundant immune cell types in the leptomeninges that contribute to cortical pathology that cannot be erased by a singular immunomodulatory drug.

To gain mechanistic insights into what drives disease progression in ageing MS patients, an animal model that replicates non-remitting clinical disability that is accompanied by unrelenting grey matter demyelination and neurodegeneration is required. We have previously shown that experimental autoimmune encephalomyelitis (EAE) induced by adoptive transfer of encephalitogenic Th17 cells into young (6-weeks-old) SJL/J recipient mice results in stromal cell remodelling within the brain leptomeninges that is accompanied by chemokine and cytokine expression. This stromal cell remodelling creates an immunocompetent niche in the leptomeninges^23^ that is spatially associated with subpial cortical grey matter demyelination, microglial/macrophage activation, disruption of the glial limitans, and evidence of an oxidative stress response^24^. However, there was no evidence of synapse loss nor axonal damage in these mice, and grey matter pathology was transient, with mice recovering from the initial inflammatory event ^24^.

In the present study, we tested the effect of age on the clinical outcome of adoptive/transfer (A/T) EAE by transferring young encephalitogenic Th17 into young (6-weeks-old) vs old (8 to 15-months-old) SJL/J recipient mice. The age of the recipient mice had a profound impact on clinical phenotype brain pathology. While young recipients underwent disease remission, signs of paralysis were more severe and sustained in old mice. Old mice exhibited numerous and large aggregates of immune cells in the leptomeninges overlying regions of cortical injury and brain atrophy. Single-cell RNA sequencing of leptomeningeal resident cells identified a number of gene expression changes that are unique to the aged EAE phenotype, which led us to examine the differential abundance of neutrophils and B cell in the leptomeninges of old vs young mice. Importantly, we validated the presence of neutrophils in the brains of a subset of progressive MS patients via post-mortem human brains collected at rapid autopsy. Collectively, our study provides a new model for studying Th17 cell-induced grey matter injury and sheds new light on potential drivers of age-dependent MS disease progression.

## Results

### Adoptive transfer of encephalitogenic T cells into old SJL/J mice results in non-remitting EAE which is independent of vivaria and sex

We have previously shown that A/T of encephalitogenic Th17 cells into SJL/J recipient mice induces EAE, with clinical symptoms first observed at approximately 5 days post-A/T, peaking at approximately day 11-12^23,24^, and recovering shortly thereafter around day 14 (**Fig. 1A**). Since age is the strongest predictor of progression^25,26^, we tested whether transfer of young Th17 cells into old recipient mice would alter the clinical course of EAE (**Fig. 1A**). While we found that old female mice (8 months) that received PLP-primed Th17 cells from young mice exhibited similar peak disease as young recipient mice, the old mice failed to remit, sustaining disability with average clinical scores of 11-13. Middle-aged mice (6 months) displayed an intermediate phenotype with some mice experiencing remission and others exhibiting non-remitting disease suggesting that 6 months of age appears is an inflection point between remission and non-remission phenotypes (**Fig. 1B-C**). Disease in both young and old mice was specific to encephalitogenic PLP139-151-primed T cells since transfer of cells from donors immunized with OVA326-339 failed to induce EAE (**Supp. Fig. 1A-B**). We also confirmed that the old vs young phenotype could be replicated in a different vivaria (University of British Columbia (UBC) and University of Toronto (UofT)) (**Fig. 1**), suggesting that age – not housing conditions – is the main driver of this clinical phenotype. Lastly, we found that the non-remitting disease course was reproduced in older mice up to 15 months of age with survival rate decreasing with increasing age (**Supp. Fig. 1**); was reproduced in male recipient mice (**Fig. 1D-E**); and persisted over several months (**Fig. 1E**).

**Figure 1.**
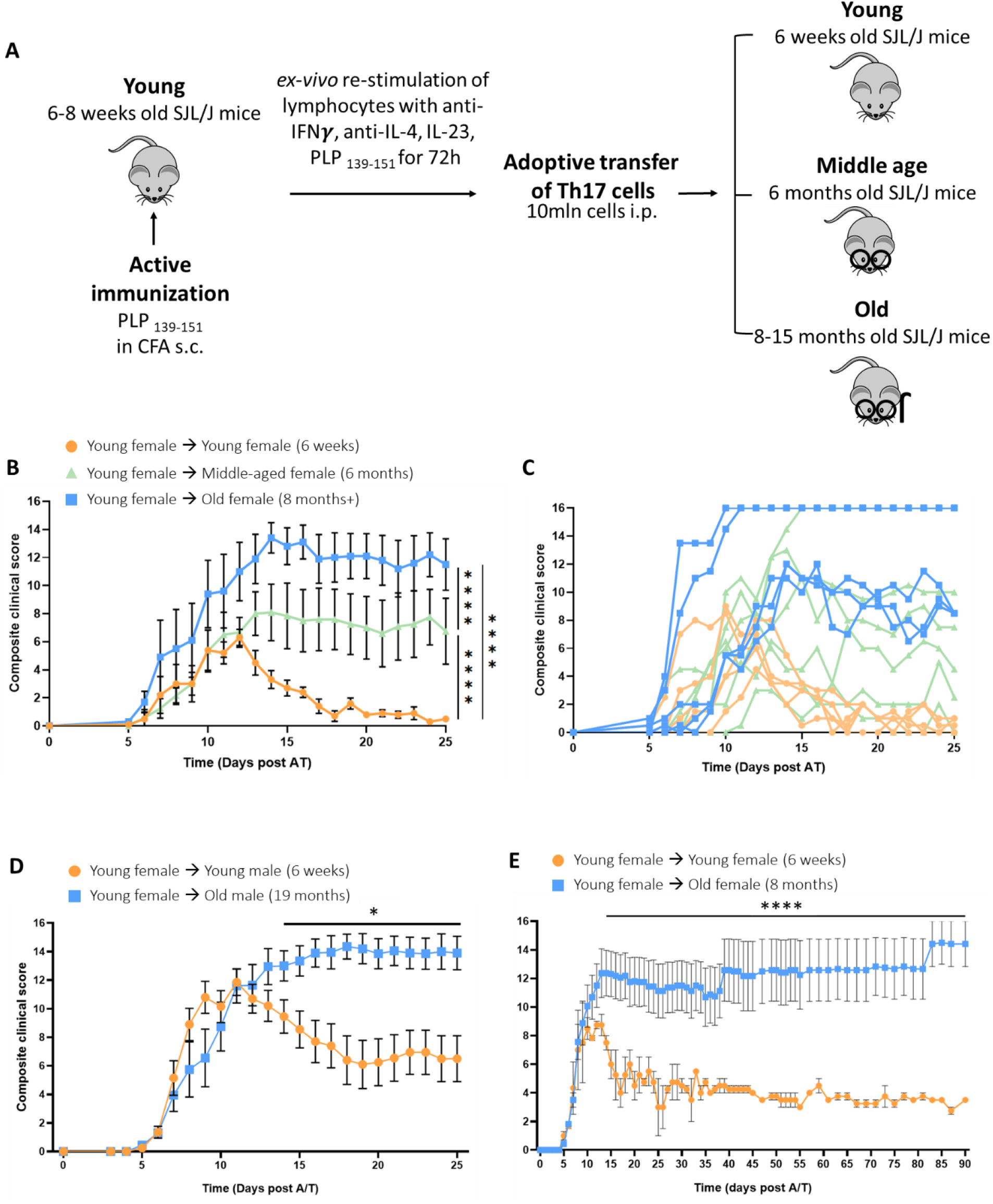
Ageing determines clinical course in SJL/J A/T EAE mice and is independent of sex, vivaria, and persists over multiple months. **(A)** Induction of adoptive transfer EAE in SJL/J mice. **(B-C)** Compositive and individual clinical scores (16-point scale) of young, middle aged, and old female A/T SJL/J EAE mice receiving cells from young female donors primed with PLP139-151. **(D)** Composite clinical score of young and old male A/T SJL/J EAE mice receiving cells from young female donors primed with PLP139-151. **(E)** Composite clinical score of young and old female SJL/J A/T EAE mice followed for up to 90 days post-adoptive transfer. Stats performed by two-way ANOVA with Bonferroni correction for multiple comparison. Error bars indicate mean with SEM. Experiments in **(B, C)** were performed at UBC while experiments in **(D, E)** were performed at U of T.

### A/T EAE in old SJL/J mice exhibit grey matter pathology reminiscent of progressive MS

We have previously shown that A/T of encephalitogenic Th17 cells into young SJL/J recipient mice provokes the formation of leptomeningeal immune cell aggregates overlaying areas of subpial demyelination at the acute phase of disease, particularly in proximity of the cortex, hippocampal fissure and brainstem (**Fig. 2A**)^23^. Histological analysis of brain tissue at day 25 post-A/T, a timepoint when young mice have largely remitted in terms of their clinical scores, revealed larger and more numerous aggregates of immune infiltrates in the brain leptomeninges in old recipients compared to young recipients. This was observed in areas of the leptomeninges proximal to the cortex (**Fig. 2B, E**), the hippocampus (**Fig. 2C, F**), and the brainstem (**Fig. 2D, G**). Quantification of imaging data revealed a 2-fold increase in the number of leptomeningeal aggregates (mean old = 3.0 TLT / mouse, mean young = 1.2 TLT/ mouse, *p* = 0.0059) and 1.5-fold increase in aggregate area in old mice compared to young (mean old = 0.024mm^2^/TLT, mean young = 0.016mm^2^/TLT, *p* = 0.0358) (**Fig. 2H, I**). Further examination of these aggregates by immunofluorescence (IF) revealed that, consistent with our previous observations^24^, leptomeningeal aggregates contained CD3^+^ T cells and B220^+^ B cells (**Fig. 2K-M, O-Q**), and were associated with a network of fibronectin^+^ extracellular matrix (**Fig.2J, N)**.

**Figure 2.**
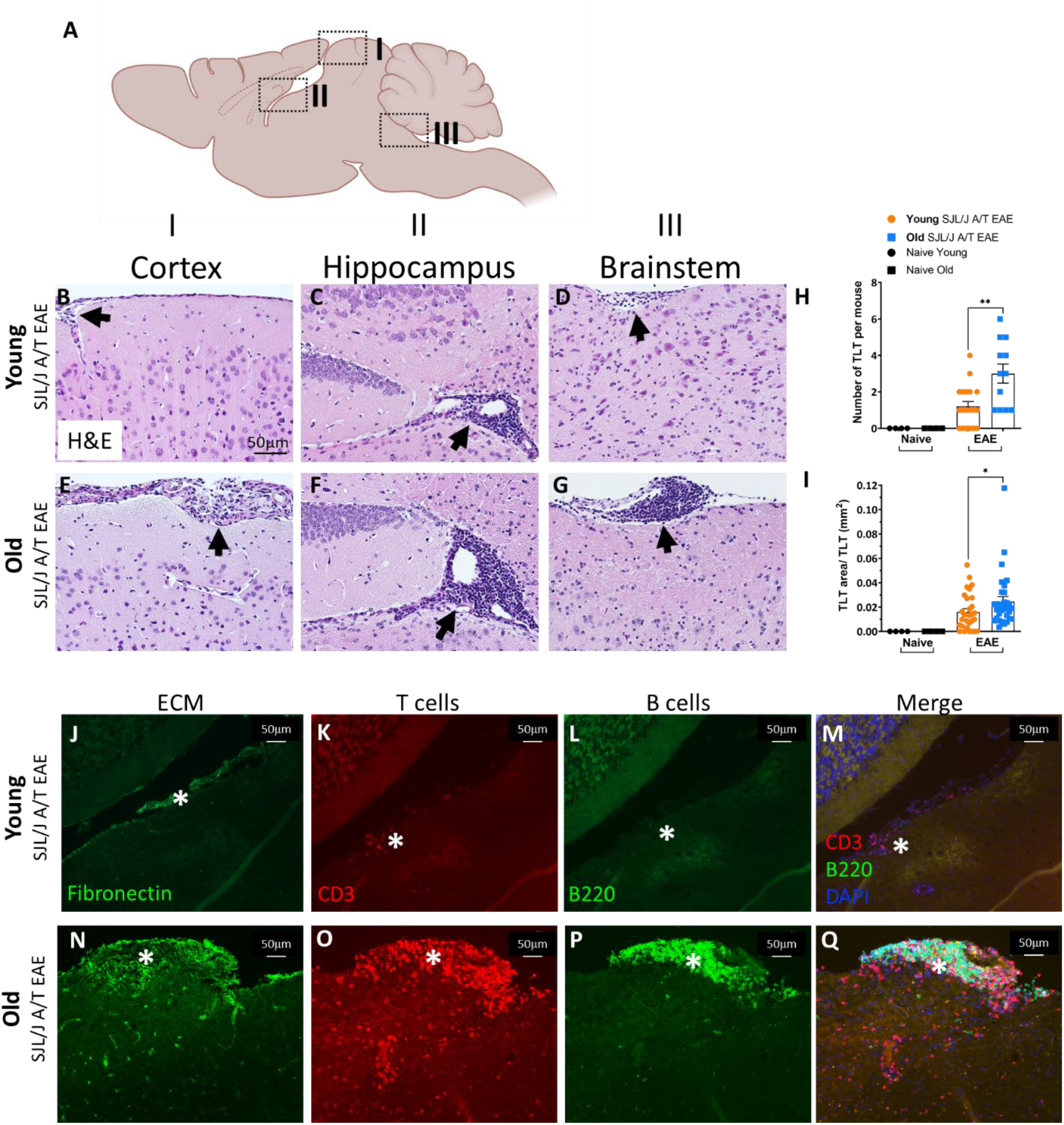
Ageing induces accumulation of lymphocytes in meninges adjacent to subpial and periventricular areas in SJL/J A/T EAE mice at post-acute disease stage. **(A)** Schematic of mouse brain with the sagittal plane in view showing (I) cortex, (II) hippocampus and lateral ventricle, and (III) brainstem and 4th ventricle regions. Representative sagittal sections of brains obtained from **(B-D)** young vs **(E-G)** old SJL/J A/T EAE mice at day 25 post-adoptive transfer. Brains of old mice show infiltration of cells in the meninges in all three locations denoted by black arrows. **(H)** Quantification of number of aggregations as well as **(I)** area covered by meningeal aggregations from old vs young mice. **(J, N)** Immunofluorescence staining for fibronectin (extracellular matrix) revealing an elaborated fibronectin+ ECM network (denoted with *). Staining of serial sections for **(K, O)** CD3, and **(L, P)** B220 reveals CD3+ T cells and B220+ B cells present within this fibronectin+ meningeal niche **(M, Q)**. Error bars indicate mean with SD.

To ascertain the impact of SJL/J A/T EAE on the grey matter in old vs young recipient mice, we performed immunohistochemistry (IHC) for proteolipid protein (PLP, myelin), glial fibrillary acidic protein (GFAP, astrocytes), Ionized calcium-binding adaptor protein-1 (Iba-1, microglia/macrophages), neurofilament (axons) and synaptophysin (synapses). At day 25 post A/T, we observed a 3-fold increase in demyelinated subpial area (mean old = 4.6% PLP^+^ area, mean young = 13.6%, *p* < 0.0001) (**Fig. 3A-C**), enhanced disruption of the glial limitans (mean old = 41.4% uninterrupted glial limitans, mean young = 86.3%) (**Fig. 3D-F**), a 2-fold increase in density of microglia/macrophage (mean old = 671 Iba-1^+^ cells/mm^2^, mean young = 375 Iba-1^+^ cells/mm^2^, *p* = 0.0165) (**Fig. 3G-I**), a 2-fold decrease in axonal integrity (mean old = 11.3% pan-neurofilament^+^ area, mean young = 20.8%, *p* = 0.0009) (**Fig. 3J-L**), and a 1.5-fold decrease in synaptic density (mean old = 35.3% synaptophysin^+^ area, mean young = 53.8% synaptophysin^+^ area, *p* = 0.0002) (**Fig. 3M-O**) in brain regions adjacent to the leptomeningeal aggregates of immune cells in old vs young SJL/J A/T EAE mice. These data indicate that old mice exhibit more demyelination, impaired glial limitans integrity, accumulation of microglia/macrophages, and more axonal and synaptic loss in the cortex compared to young mice at day 25 post-A/T.

**Figure 3.**
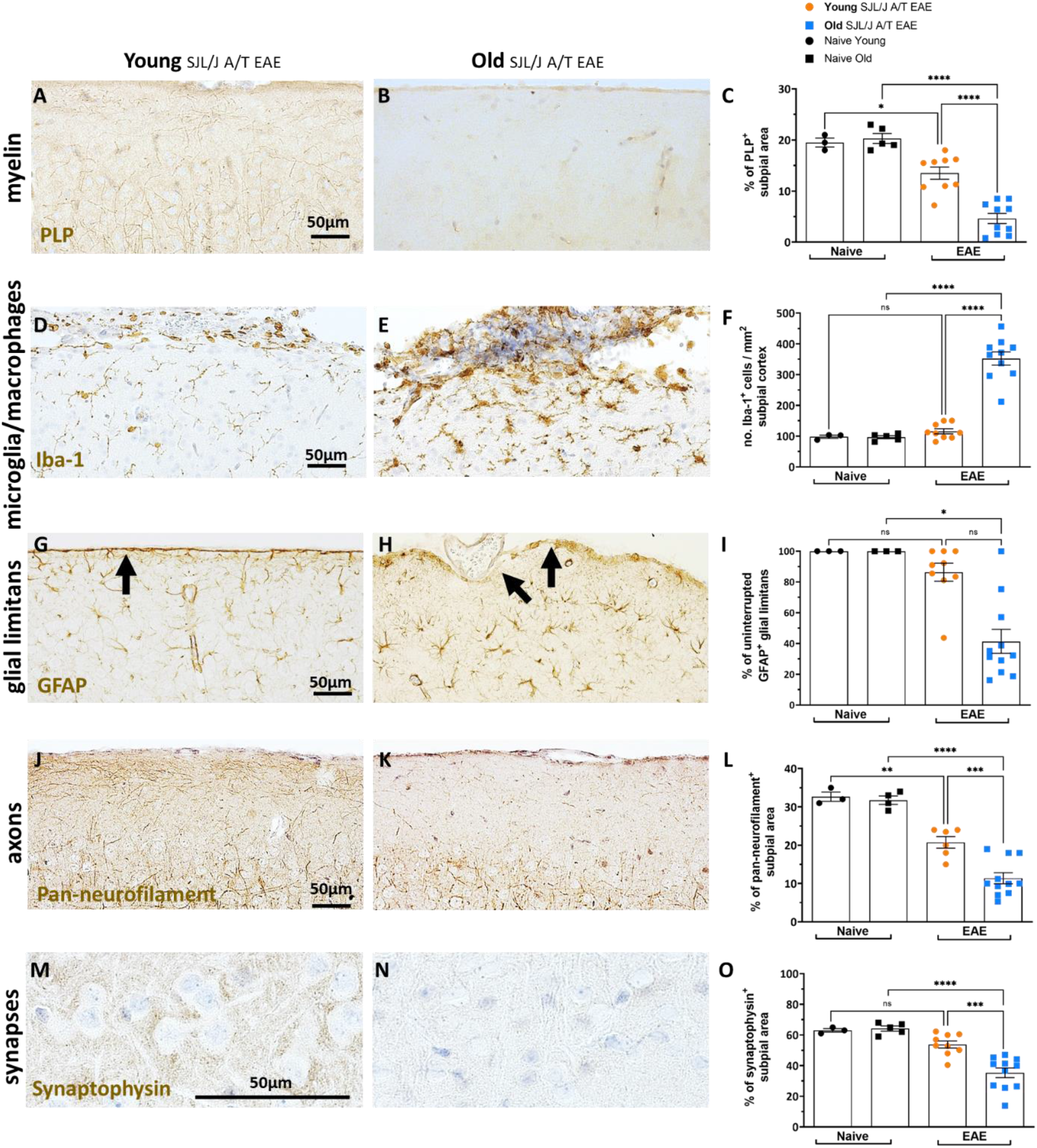
Ageing induces sustained subpial cortical injury in old SJL/J A/T EAE mice. Assessment of the subpial somatosensory cortex of old vs young SJL/J A/T EAE mice at the post-acute stage (D25 post A/T) by immunohistochemistry (IHC). **(A-C)** Myelin content (PLP); **(D-F)** Macrophage/microglia density (Iba-1); **(G-I)** Glial limitans and astrocytes (GFAP); **(J-L)** axons (pan-neurofilament). Statistics C, F, L = One-way ANOVA with Tukey’s correction for multiple comparisons. Statistics I = Kruskal Wallis with Dunn’s correction for multiple comparisons. Error bars indicate mean with SD.

Serum neurofilament light chain (sNfL) has become an increasingly utilized biomarker for ascertaining MS severity^27^. To test whether sNfL is induced by A/T of Th17 cells, we subjected serum from old vs young SJL/J A/T EAE mice to Quanterix single-molecule array (Simoa) technology. At peak disease, old and young mice exhibited similar levels of serum NfL (**Supp. Fig. 2A**). However, at the post-acute phase of disease, old recipients exhibited augmented sNfL compared to sex-matched young recipients (mean old = 2384 pg/mL, mean young = 1123 pg/mL) (**Supp. Fig. 2B**), and sNfL levels positively correlated with disease severity (Spearman’s r = 0.5053, *p* = 0.0273) (**Supp. Fig. 2C**). To determine whether sNfL levels reflect ongoing neuronal damage in the subpial cortex, we assessed brain tissues from corresponding mice by IHC. Old mice exhibited lower %NfL^+^ area in the subpial cortex compared to young mice (mean EAE old = 11.0% NfL^+^ area, mean young = 17.5% NfL^+^ area, *p* < 0.0001) (**Supp. Fig. 2E**), and these values negatively correlated with sNfL levels (Spearman’s r = - 0.7818, *p* = 0.0105) (**Supp. Fig. 2F**), suggesting that increased sNfL levels reflect a decrease in NfL in the cortex. Therefore sNfL levels correlate with worse clinical outcomes in old mice, and elevated sNfL in old mice is driven at least in part by neuronal damage in the cortex.

A key hallmark of progressive MS is a reduction in brain volume driven in part by atrophy in the cortical grey matter^28,29^. To test if the SJL/J A/T EAE model in old mice recapitulates brain atrophy, we followed old and young SJL/J mice for 90 days post-A/T and assessed brain volume by T2-weighted 7-Tesla magnetic resonance imaging (MRI) at three timepoints – acute (Day 11), post-acute 1 (Day 39), and post-acute 2 (Day 90) (**Supp. Fig. 2G**). We found that old SJL/J A/T EAE mice exhibited lower brain volume compared to age- and sex-matched controls at both the post-acute timepoints, while young EAE mice exhibited similar brain volume compared to their appropriate controls (**Supp. Fig. 2H**). Moreover, when examining each brain region separately, decreased brain volume in old SJL/J A/T EAE mice was most pronounced in the somatosensory cortex at the second post-acute timepoint (**Supp. Fig. 2I**). Although we did not have enough mice to establish significance, these data suggest that prolonged EAE may result in diminished brain volume in old SJL/J A/T EAE mice, detectable by MRI as early as D39 post A/T.

### Ageing does not impact CNS-resident T cells in SJL/J A/T EAE

The divergent clinical and pathological (grey matter) phenotype in old vs young SJL/J A/T EAE model emerges after the peak of disease. Thus, examining the composition of immune cells in the leptomeninges at peak disease provides an opportunity to assess what may be responsible for the subsequent poor outcome in old recipient mice. We first asked whether old vs young mice differed in the composition and phenotype of T cells at peak disease by performing flow cytometry on whole brains and spinal cords. We found no differences in the frequencies and absolute numbers of CD4^+^/CD8^+^ T cells derived from the brain and spinal cord of old vs young SJL/J A/T EAE mice (**Fig. 4A, B**). Furthermore, *ex vivo* restimulation of T cells from whole brains and spinal cords of old vs young SJL/J A/T EAE mice taken at the acute time point revealed no difference in their capacity to produce IL-17, GM-CSF, and IFNγ (**Fig. 4C, D**). These data show that differences in CNS resident T cells are not likely to account for altered clinical and pathological attributes of SJL/J A/T EAE in old vs young mice.

**Figure 4.**
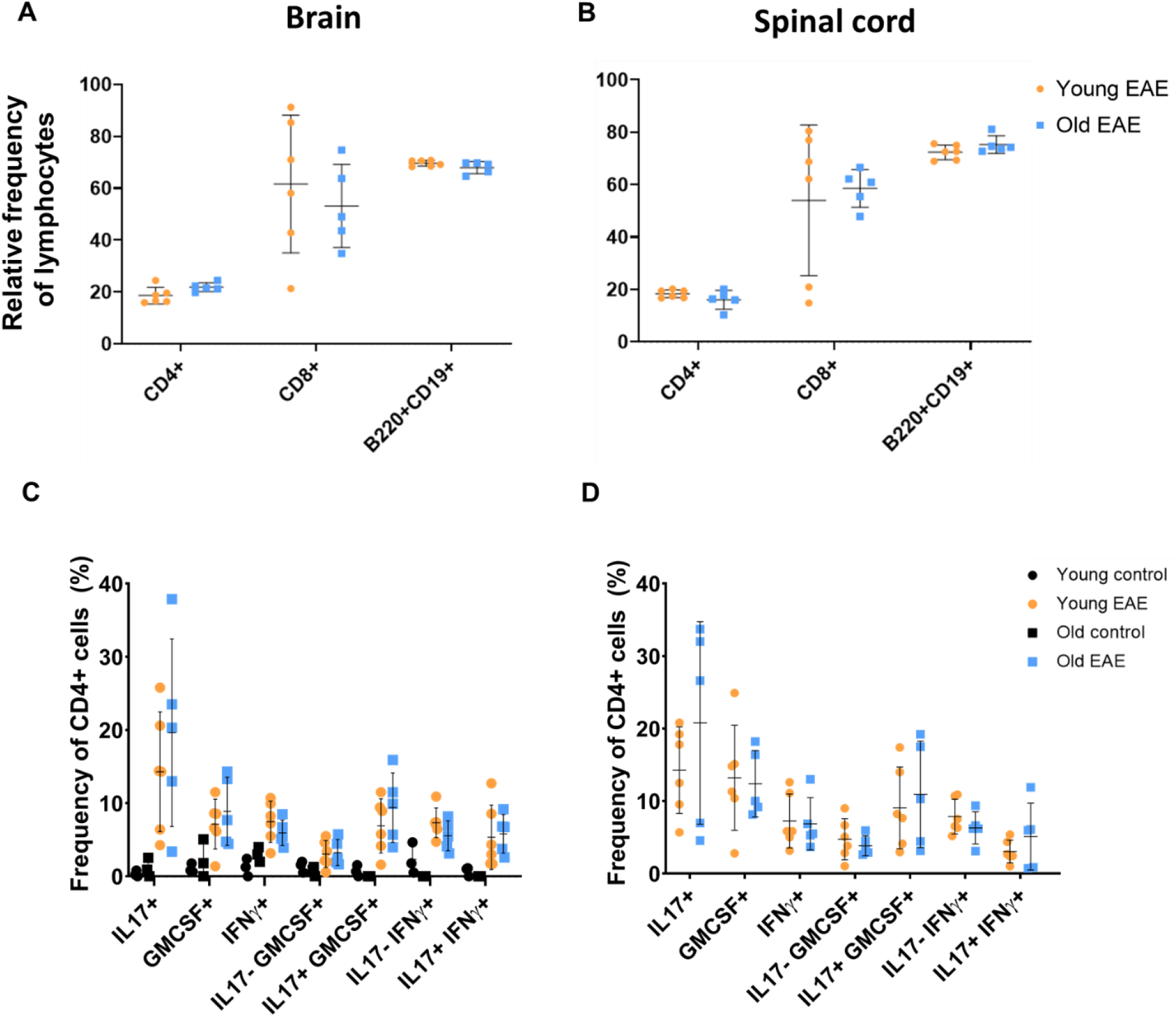
Ageing does not impact the frequency of lymphocytes or the ability of T cells to produce cytokines in SJL/J A/T EAE mice at peak disease stage. Whole brains and spinal cords of young and old SJL/J A/T EAE and naive mice were analyzed by flow cytometry. **(A-B)** Frequencies of B220+CD19+ B cells, CD4+ and CD8+ T cells of all CD45+ cells in the brains and spinal cords of old vs young SJL/J A/T EAE mice. Stats by Mann-Whitney U showed no statistical difference between the groups. Cells were also stimulated with PMA and ionomycin for 5 hours. **(C-D)** Frequencies and absolute numbers of cytokine-producing CD4+ T cells was assessed using intra-cellular staining for GM-CSF, IFNγ, and IL-17. Statistical test by Mann-Whitney showed no significant differences between the groups. Error bars indicate mean with SD.

### Old SJL/J A/T EAE mice exhibit a deficit of B cells and monocytes and an accumulation of neutrophils in the leptomeninges

Using the whole brain to ascertain the impact of age on T cell phenotype in the context of SJL/J A/T EAE may have obscured brain compartment-specific effects. Indeed, we know that T cells accumulate in the leptomeninges in both old and young SJL/J A/T EAE mice, and the leptomeninges represents only a fraction of the entire brain. To interrogate differences in compartment-specific populations, we applied flow cytometry to single cell suspensions released from separately dissected leptomeningeal, cortical, and brainstem brain fractions disease, focusing again on the peak acute disease timepoint prior to the bifurcation of clinical phenotypes in old *vs* young mice in order to gain insight into what may drive non-remitting EAE. Despite separating by brain region, we still found no differences in CD3^+^CD4^+^ T cell numbers or frequencies in the leptomeninges, cortex, or brainstem, confirming our earlier results on whole brain (**Fig. 5A**). However, in the leptomeninges we observed a 1.5-fold decrease in CD11b^+^Ly6C^+^ monocytes (*p* < 0.0001) (**Fig. 5B**), and a 3-fold decrease in absolute number (*p* = 0.0072) and frequency (*p* = 0.0314) of CD19^+^B220^+^ B cells in the leptomeninges of old SJL/J A/T EAE mice compared to young mice (**Fig. 5C**). We also observed a 2-fold increase in frequency of CD11b^+^Ly6G^+^ neutrophils (*p* < 0.005) (**Fig. 5D**. When examining the brain parenchyma (cortex and brainstem), we only noted a significant decrease in B cells, a slight trend toward increase in frequency and absolute number of neutrophils and no difference in frequencies or absolute number of monocytes (**Supp. Fig. 4**). Thus, alterations in monocytes, B cells and neutrophils in old *vs* young recipient mice during the acute disease timepoint is largely restricted to the leptomeninges. To further confirm an accumulation of neutrophils in the brains of old SJL/J A/T EAE mice, we performed IF, staining for Ly6G, a marker of neutrophils. Indeed, we observed an accumulation of Ly6G^+^ cells in the leptomeninges overlying the cortices (mean old = 327.6 cells, mean young = 84.8, *p* < 0.001) and brainstems (mean old = 907.2 cells, mean young = 226.7, *p* < 0.01) of old mice (**Fig. 5E-H**).

**Figure 5.**
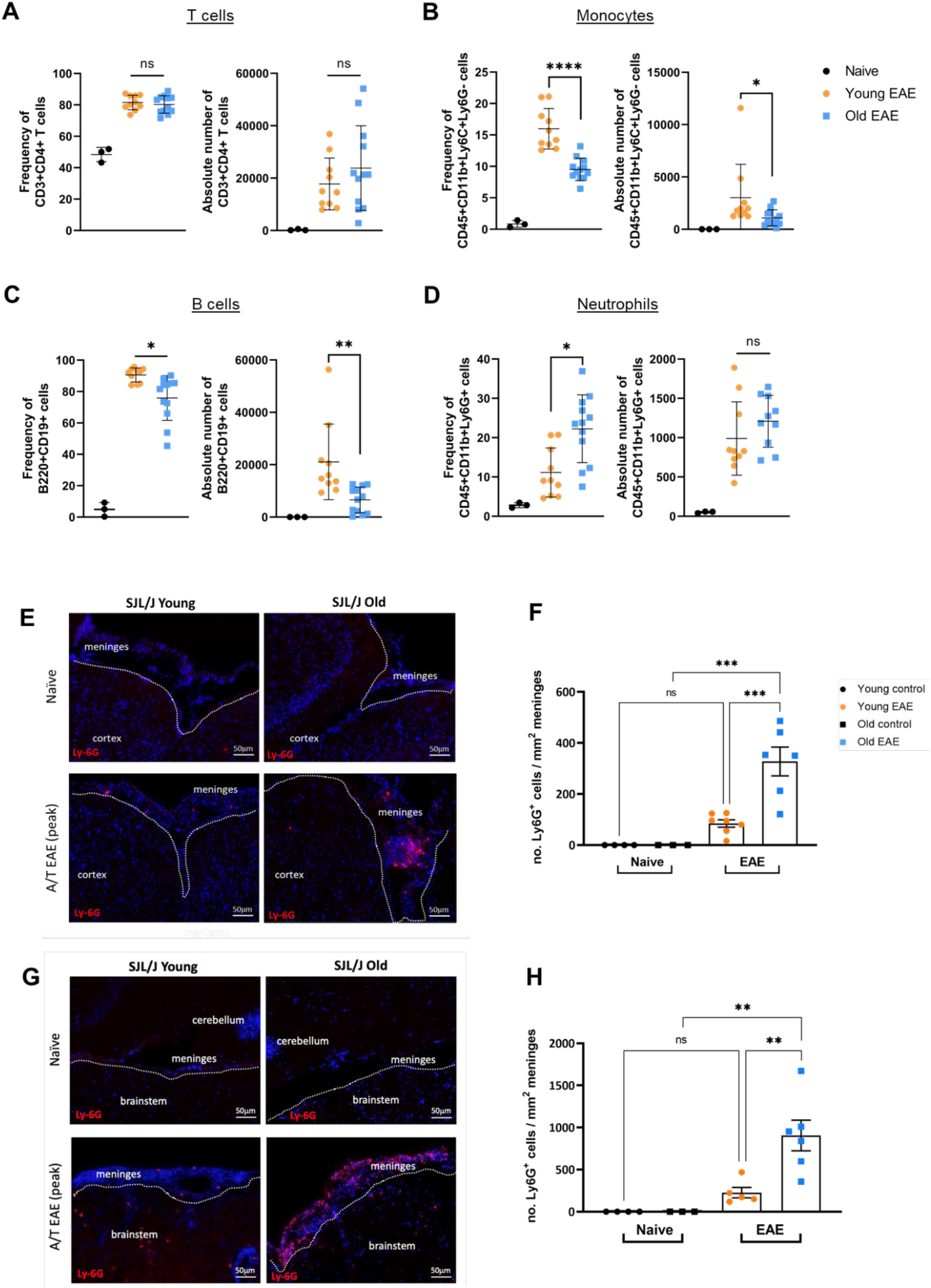
Flow cytometry and immunofluorescence of the brain from old vs young SJL/J A/T EAE mice reveals differences in immune cell compositions. Leptomeninges was separately dissected from mice at the acute timepoint of EAE and subjected to analysis by flow cytometry. Despite no change in **(A)** T cell number or frequency, we observed a decrease in density of **(B)** monocytes and **(C)** B cells, and an increase in frequency of **(D)** neutrophils. We also performed immunofluorescence staining for Ly6G, identifying neutrophils in fresh-frozen brain tissue from old vs young naïve and SJL/J A/T EAE mice. Note the enrichment in Ly6G+ neutrophils in the meninges overlaying the **(E-F)** cortex and **(G-H)** brain stem. Stats A-D = one-way ANOVA (absolute number) or Kruskal-Wallis (frequency) with correction for multiple comparisons. Stats F, H = Kruskal Wallis with correction for multiple comparisons. Data is expressed as mean with SD.

Collectively, these data demonstrate that while ageing does not impact the number or phenotype of brain-resident CD3^+^CD4^+^ T cells, there is a significant age-dependent difference in the accumulation of leptomeningeal B cells, neutrophils, and monocytes in the context of SJL/J A/T EAE.

### Transcriptomic analysis of the EAE leptomeninges reveals gene expression differences between old and young SJL/J A/T EAE mice

We next asked whether the transcriptomic landscapes of immune cells differed in the leptomeninges and cortex from old vs young mice at peak disease using single-cell RNA sequencing (scRNAseq). We accomplished this by submitting single cell suspensions derived from the leptomeninges of old *vs* young SJL/J A/T EAE mice to sequencing on the 10X Genomics platform. We obtained a total of 18,321 cells from two independent experimental repeats. After performing quality control and data integration pipelines in Seurat V3.0^30^, 15,641 cells were subjected to unsupervised UMAP clustering and 15 clusters were identified based on differential gene expression (**Fig. 6A**). We ascribed putative identities of these clusters based on dominant cluster-specific genes, including those for neutrophils (*Mmp8, Mmp9, Cxcr2, Ly6g*), macrophages/monocytes (*Itgam, Ly6c1, Ly6c*), T cells (*Cd3e, Cd4, Cd8a, Tcf7*), and B cells (*Cd19, Ighm, Ighd*) (**Fig. 6B**). We then interrogated the gene expression profiles of B cell, neutrophil, and monocyte clusters because these populations were differentially represented in old vs young mice by flow cytometry. We noted 495 differentially expressed genes among the B cell clusters, 473 genes in the neutrophil cluster, 107 genes among the monocyte/macrophage clusters and only 82 genes among the T cell clusters in old *vs* young SJL/J A/T EAE mice. We then filtered for genes with a *p*-value of <0.01. In the B cell clusters, we found that young EAE mice upregulated more transcripts involved in B cell development such as *Ighm, Ebf1, Iglc3, Cd79b, Ighd*, and *Vpreb3*, while old EAE mice upregulated pro-inflammatory genes such as *Il7r, Fos, Ccl5, Ccl17, Ighg2b, Bcl2a1b, Syngr2, Cxcl16* and *Apoe* (**Fig. 6C**). Of the genes in the neutrophil clusters, we found that young EAE mice expressed more transcripts for *C3* and *Cd74*, while old EAE mice expressed more transcripts associated with innate immunity such as *Chil3, Itgam, Il18rap, Retnlg, Cxcl2, Hmgb2*, and *Cd14* (**Fig. 6D**). Lastly, in the monocyte/macrophage clusters, we found young EAE mice had higher levels of transcripts for *Ly6i, Ccl5, C3, Chil1, Ly6c2, Cx3cr1*, and *Cxcl9*, while old EAE mice had higher levels of transcripts for inflammation and complement pathways such as *Ccl2, Tnf, Ccr1, Cd93, C1qc, C1qa, C1qb, Cd14*, and *Apoe* (**Fig. 6E**). Of the T cells, young EAE mice expressed higher levels of *Nrgn*, while old EAE mice expressed high levels of *Fos, Ctla2a*, and *Ramp3* (**Fig. 6F**). These data further suggest that while T cells are important in establishing disease and initiating pathogenesis, other cells such as B cells, neutrophils, and monocytes/macrophages contribute to the differential clinical courses between old vs young SJL/J A/T EAE mice.

**Figure 6.**
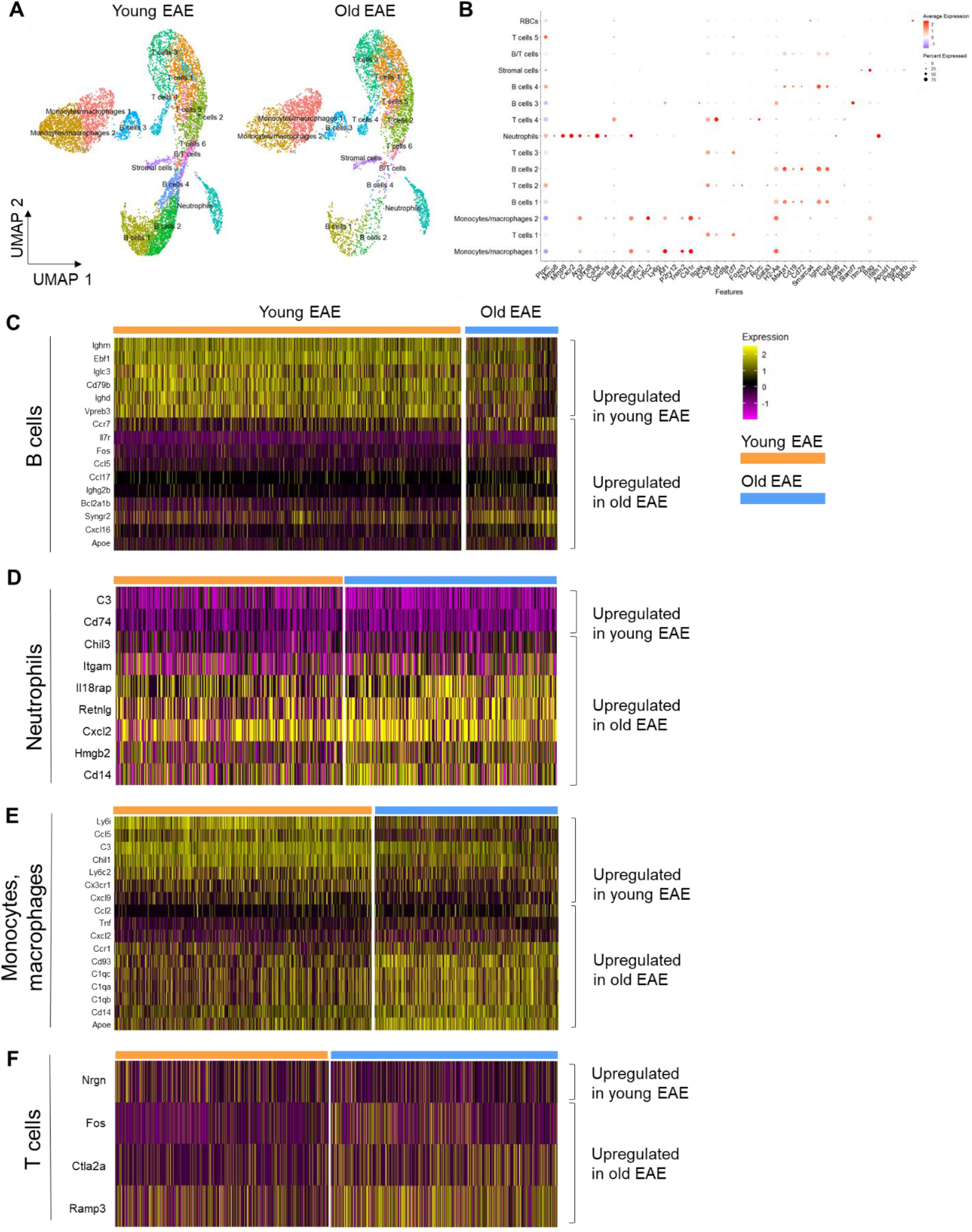
Transcriptomic analysis of the SJL/J A/T EAE meninges reveals age-dependent heterogeneity in transcriptional profiles of B cell, neutrophil, and monocyte/macrophage populations at peak disease stage. Leptomeninges of old vs young SJL/J A/T EAE mice at peak disease were dissected and sent for single-cell RNA sequencing on the 10X genomics platform. **(A)** UMAP clustering reveals proportional differences in leptomeningeal immune cell populations. **(B)** Identification of clusters was performed using transcripts for lineage-specific markers. Differential gene expression analysis based on age was performed on **(C)** B cell, **(D)** neutrophil, **(E)** monocyte/macrophage clusters and **(F)** T cells. Gene expression heatmaps were generated from select transcripts that exhibited a *p* < 0.01 and log2 fold-change cutoff of 0.5.

### Neutrophils populate the leptomeninges of progressive MS brains

Neutrophils have been implicated in demyelination and axonal degeneration during the acute phase of EAE in in C57BL/6 mice^31^, but little is known about their presence in the MS brain. We therefore examined previously characterized post-mortem MS brains, which showed a range of subpial demyelination and meningeal inflammation compared to age-matched non neurological controls^32^ for the presence of neutrophils in the leptomeningeal compartment using high magnification microscopy on H&E-stained samples. Neutrophils were identified based on their multi-lobular nuclei. While rare, we found neutrophils in the leptomeninges of a subset of MS donors (**Fig. 7A**). We then stratified donors based on the presence or absence of neutrophils in the leptomeninges and performed correlation studies comparing the extent of subpial demyelination and leptomeningeal inflammation with the presence or absence of neutrophils. The donors with leptomeningeal neutrophils exhibited a higher % of demyelination in the subpial cortex compared to donors without leptomeningeal neutrophils (63.8% vs 41.3% demyelination, *p* < 0.05) (**Fig. 7B**), as well as a higher number of leptomeningeal CD20^+^ B cells (10.0 CD20^+^ cells/mm length of leptomeninges vs 6.1 CD20^+^ cells/mm length of leptomeninges) (**Fig. 7C**). While we did not find a significant association between the presence of neutrophils and the number of leptomeningeal CD3^+^ T cells/mm length of leptomeninges in these donors, a trend toward more CD3^+^ T cells in the leptomeninges of patients with neutrophils than those without was observed (**Fig. 7D**). In conclusion, we identified neutrophil accumulation in the leptomeninges of a subset of progressive MS patients which also exhibit leptomeningeal inflammation and cortical subpial demyelination.

**Figure 7.**
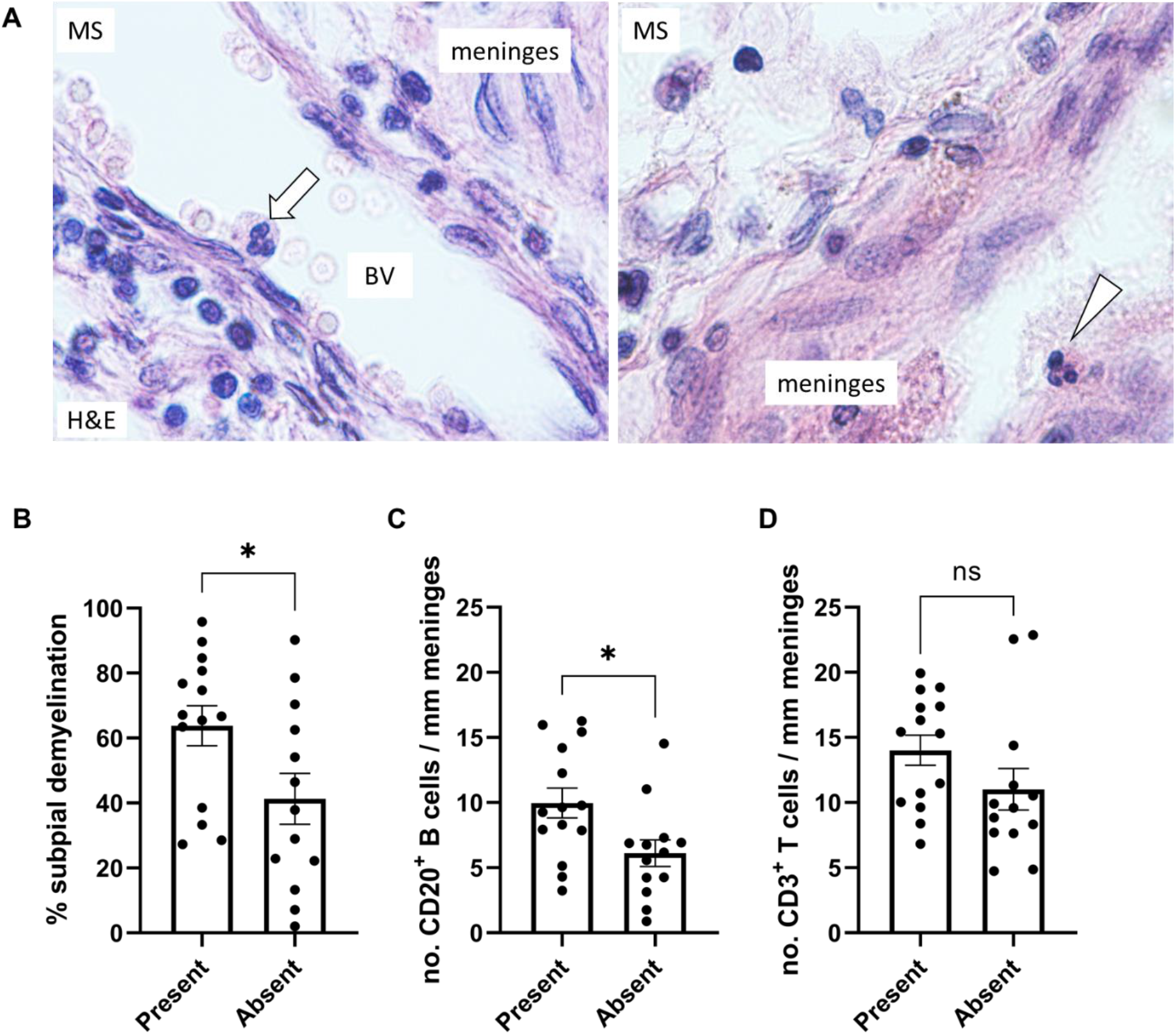
The presence of neutrophils in the meninges of patients with progressive MS associates with more extensive subpial demyelination and a higher density of meningeal B cells. **(A)** Representative hematoxylin and eosin (H&E) staining of formalin-fixed paraffin-embedded cortex from progressive MS patients (n=27) showing a neutrophil localized within a blood vessel (BV) (arrow) and a neutrophil localized outside of blood vessels (arrowhead) in the meninges lining the cortex. Quantification of **(B)** % subpial demyelination, **(C)** density of meningeal CD20+ B cells and **(D)** density of meningeal CD3+ T cells in MS donors with or without neutrophils identified outside BV in the leptomeninges. Stats by unpaired student’s t-test, * indicates *p* < 0.05.

## Discussion

In this study, we show that adoptive transfer of PLP-primed encephalitogenic Th17 cells into old SJL/J mice induces non-remitting EAE. Furthermore, the non-remitting clinical course was accompanied by leptomeningeal inflammation, grey matter demyelination, axonal damage, synapse loss and disruption of the glial limitans, as well as brain atrophy and accumulation of neurofilament light chain in the serum, all hallmarks of progressive MS. We also show that this model is highly reproducible across vivaria and is independent of sex of recipient mice, providing further evidence that age is the primary driver of the clinical and pathological phenotype. Therefore A/T of encephalitogenic Th17 cells into old *vs* young SJL/J mice is a valuable method for ascertaining the role of ageing on grey matter pathology associated with leptomeningeal inflammation in EAE in the absence of the confounding adjuvant driven/cytokine-storm effects that occur with active EAE or lentiviral-based introduction of cytokines^31,33,34^.

We have previously shown that cytokines such as IL-17A, IL-17F, IL-22 and Lymphotoxin exert direct effects on the underlying stromal cells of the leptomeningeal subarachnoid space, resulting in the release of chemokines and cytokines that in turn recruit and tune additional immune cells (T cells, B cells, myeloid cells etc) that infiltrate the leptomeninges. This results in the formation of so-called tertiary lymphoid tissues (TLT) that exhibit varying degrees of complexity and organization^23^. Although these structures are also present in early MS^9^, they may play a particularly pathogenic role in the progressive phase of the disease^20,35^. Thus, one potential reason for inefficacy of immunomodulatory therapies in PMS^36^ could be due to their lack of access to these structures and/or redundant immune-mediated mechanisms that sustain these structures which cannot be silenced by a single therapy. Since these structures are evident from the very earliest stages of MS, we reason that they may be fueling a form of “silent progression” that ultimately impacts the health of the underlying grey matter.

While age is the strongest predictor of MS progression^25^, we do not know what aspects of ageing impact the MS brain such that it becomes more susceptible to grey matter injury. One possible explanation for the severe EAE phenotype in old SJL/J recipient mice is that young Th17 cells become more activated in the environment of the old CNS. However, we found no difference in the quantity or quality of CNS-resident CD4^+^ T cells, suggesting that other age-associated changes are dominant factors that determine clinical outcome. Instead we noted a paucity of monocytes/B cells and an increase in neutrophils in the leptomeninges overlying the cortex and brainstem of old SJL/J A/T EAE mice compared to young SJL/J A/T EAE mice by flow cytometry and immunofluorescence microscopy. These experiments were performed at the peak of disease, prior to the bifurcation of the old *vs* young clinical phenotype. Thus, changes at the acute phase may “set up” the CNS for post-acute clinical disease.

To gain insight into the functional states of each cellular compartment associated with the remitting or non-remitting disease course in SJL/J A/T EAE mice, we chose a single-cell transcriptomic approach. We interrogated differences in gene expression of leptomeningeal T cells, B cells, neutrophils, and monocyte/macrophages of old vs young SJL/J A/T EAE mice at peak disease. While T cell clusters showed a similar transcriptional signature, the B cells lineage showed an upregulation of several genes associated with developing B cells in young mice (*Ebf1, Vpreb3, Cd79b, Ighm, Ighd*), while more genes associated with mature and inflammatory B cells were upregulated in old mice (*Ighg2b, Bcl2a1b, Fos, Cxcl16, Ccl5, Ccr7*). Within the neutrophil cluster, we found an upregulation of genes associated with phagocytosis and immune activation (*Cd14, Itgam, Hmgb2*) as well as an upregulation of transcripts for the neutrophil chemoattractant *Cxcl2*, which has been reported in the literature to be expressed by activated neutrophils to facilitate recruitment of more neutrophils^37,38^. In the monocyte/macrophage clusters, we found cells from old mice exhibited an upregulation of transcripts for inflammatory cytokines (*Cxcl2, Tnf, Ccl2*) and phagocytosis (*Cd14, Apoe*) as well as the classical complement pathway (*C1qa, C1qb, C1qc, Cd93*). The latter is known to be involved in the stripping of synapses^39,40^ and could therefore be a potential mechanism at play in the brain of old EAE mice where we observed a loss of synapses in the cortex.

An important question is whether our model is generalizable across genetic backgrounds, and more broadly relevant to MS disease progression. Although C57BL/6 mice exhibit predominantly spinal cord pathology, Segal and colleagues have recently shown that age is associated in that strain with a non-remitting phenotype that bears many similarities to what we observe here in SJL/J mice (personal communication), suggesting that age is a powerful modifier of CNS pathology independent of strain differences. In comparing this model to human MS, not only are many features of progressive MS recapitulated, but aged A/T EAE SJL/J EAE mice demonstrated leptomeningeal accumulation of neutrophils, a finding we validated in the leptomeninges of a subset of SPMS patients that also exhibited leptomeningeal inflammation and cortical demyelination. Neutrophils have likely been underestimated in MS studies, as they are innate immune cells, typically short-lived without antigen-presenting capacity, and relatively scarce in MS post-mortem brain tissues with long-standing disease^41^. However, neutrophils may contribute to MS and EAE pathogenesis in several ways. Yamasaki *et al* have shown that in the C57BL/6 model of EAE, neutrophils are capable of engulfing myelin and contributing to demyelination^42^. Neutrophils also secrete a repertoire of inflammatory mediators, including IL-1β^43,44^, which may stimulate differentiation of CD4^+^ T cells into Th17 cells. Neutrophils also produce matrix metalloproteinases (MMPs) and myeloperoxidases (MPO), which may contribute to BBB leakage and breakdown^45,46^. Indeed, in our scRNAseq data, we detected transcripts for *Mmp8* and *Mmp9*, as well as *Il1b* in the leptomeningeal neutrophil cell cluster, suggesting that neutrophils in our SJL/J A/T EAE model may be involved in mediating stromal remodeling and inflammation. It is important to note that while our EAE studies showed an enrichment of neutrophils and a paucity of CD20+ B cells in the brain leptomeninges at peak disease, the MS cohort analysed showed that donors with neutrophils in the meningeas had an enrichment of CD20 + lymphocytes in the leptomeninges. This apparent discrepancy may be due to the fact that the human tissue analysed in this study is from a cohort of progressive MS patients with long-standing disease (>20 years) as opposed to the EAE model which captures acute inflammatory changes at peak disease. Indeed, although present, the neutrophils in the progressive MS post-mortem brains were rare and B cells showed a diffuse localization throughout the leptomeninges as described in other donor cohorts^32^.

Our study has some limitations that can be followed up in the future. Specifically, because of tissue processing biases, our scRNAseq and flow cytometry data are enriched for immune cells rather than stromal or glial cell populations. We have previously shown that stromal cells in the sub-arachnoid space are dramatically remodelled following introduction of PLP-primed Th17 cells^23^, thus a better understanding of alterations in stromal cell gene expression in response to Th17 cell infiltration may provide clues into what drives sub-pial grey matter pathology. Moreover, during ageing microglia and astrocytes become more inflammatory, and neurons are increasingly prone to damage^26,47–49^. Nevertheless, the connection between inflammatory factors that originate from the leptomeninges and their impact on underlying glial cells remains relatively unexplored, and our model will serve as a valuable tool for interrogating the effects of ageing on grey matter pathology.

## Methods

### Mice

Female 6- to 10-week-old SJL/J CD45.1+ mice were obtained from Envigo. Animals were housed at the University of Toronto animal facilities under specific pathogen–free conditions, in a closed caging system with a 12-hour light/12-hour dark cycle. They were provided with a standard irradiated chow diet (Teklad; Envigo, 2918) and acidified water (reverse osmosis and ultraviolet sterilized) *ad libitum*.

At the University of British Columbia, SJL/J mice (Envigo) were bred and housed under specific pathogen-free conditions at the Center for Disease Modeling. Up to 5 mice per cage were housed in Ehret cages with BetaChip bedding and had *ad libitum* access to standard irradiated chow (PicoLab Diet 5053) and reverse osmosis/chlorinated (2-3ppm)-purifed water. Housing rooms were maintained on a 14-hour light/10-hour dark cycle with temperature and humidity ranges of 20-22° and 40-70%, respectively. All experiments were performed according to guidelines from UBC Animal Care Committee and Biosafety Committee-approved protocols.

### Induction of EAE and clinical evaluation

Donor 6-week SJL/J female mice were immunized with 100μg of PLP139-151 (HSLGKWLGHPDKF; Canpeptide) in an emulsion of incomplete Freund’s adjuvant (BD Difco), supplemented with 200 μg Mycobacterium tuberculosis H37 Ra (BD Difco, 231141) in a total volume of 300 μL administered as three 100 μL subcutaneous injections on the back and flanks. Nine days post-immunization, donors were sacrificed by CO2 asphyxiation. Subsequently, cells from spleens and lymph nodes (inguinal, axillary, brachial, and cervical) were released and cells were then restimulated *ex vivo* with PLP139– 151 (10 μg/mL) in the presence of anti–IFN-γ (20 μg/mL, Bioceros), anti–IL-4 (20 μg/mL, Bioceros), and IL-23 (10 ng/mL, R&D Systems) for 72 hours at 37°C. In total, 1 × 10^7^ cells were injected intraperitoneally into SJL/J recipient mice.

For clinical assessment of EAE disease, recipient mice were weighed daily and scored according to a composite scale that we and others have previously published. Briefly, the composite scale measures mobility impairments in each limb and the tail. Each limb is graded from 0 (asymptomatic) to 3 (complete paralysis), and the tail is graded from 0 (asymptomatic) to 2 (limp tail). Assessment of the righting reflex is scored from 0 to 2, with 0 being assigned for a normal righting reflex, 1 for slow righting reflex, and 2 for a delay of more than 5 s in the righting reflex. Each criterion was measured in 0.5 increments. Thus, the composite score ranges from 0 (nonsymptomatic) to 16 (fully quadriplegic mouse with limp tail and significantly delayed righting reflex)^50–52^.

### Histology and immunostaining

Seven-micron paraffin sagittal sections of mouse brain were collected from the midline, mounted on Superfrost Plus glass slides (Knittel Glass), and dried in the oven (Precision compact oven, Thermo Fisher Scientific) overnight at 37°C. Paraffin sections were deparaffinated in xylene (Fisher Chemical, Thermo Fisher Scientific) and rehydrated through a series of ethanol washes. Histology was performed using standard HE stain and then placed in xylene before being coverslipped with Entellan mounting media (MilliporeSigma).

For immunohistochemistry, formalin-fixed, paraffin-embedded (FFPE) slides were deparaffinized and rehydrated as described above. Slides were subsequently incubated in 0.3% H2O2 in methanol for 20 minutes to block endogenous peroxidase activity. Epitopes were exposed by heat-induced antigen retrieval in 10 mM Tris + 1 mM EDTA (pH 9.0), depending on the antibody used (see **Supp. Table 1**) in a pressure cooker placed inside a microwave set at high power (~800 watts) for 20 minutes.. Endogenous peroxidases activity was blocked by incubation in PBS with 0.3% H2O2 for 20 min at room temperature. Non-specific protein binding was blocked by incubation with 10% normal goat serum (DAKO). The nonspecific binding of antibodies was blocked using 10% normal goat serum (DAKO) in PBS for 20 minutes at room temperature. Myelin protein was detected using an antibody for proteolipid protein (PLP), microglia/macrophages were detected using an antibody for ionized calcium binding adaptor molecule 1 (Iba-1), the glial limitans was detected using an antibody for glial fibrillary acidic protein (GFAP), neurofilament was detected using an antibody for pan-neurofilament, and synapses were detected using an antibody against the neuroendocrine secretory granule membrane (Synaptophysin). Primary antibodies were applied overnight at 4°C, diluted in normal antibody diluent (Immunologic, Duiven, The Netherlands). The following day, sections were incubated with a post-antibody blocking solution for monoclonal antibodies (Immunologic) diluted 1:1 in PBS for 15 min at RT. Detection was performed by incubating tissue sections in secondary Poly-HRP (horseradish peroxidase)-goat anti-mouse/rabbit/rat IgG (Immunologic) antibodies diluted 1:1 in PBS for 30 min at RT followed by application of DAB (3,3-diaminobenzidine tetrahydrochloride (Vector Laboratories, Burlingame, CA, U.S.A.) as a chromogen. Counterstaining was performed with hematoxylin (Sigma-Aldrich) for 10 min. The sections were subsequently dehydrated through a series of ethyl alcohol solutions and then placed in xylene before being coverslipped with Entellan mounting media (Sigma Aldrich). The colorimetric staining was visualized under a light microscope (Axioscope, Zeiss), connected to a digital camera (AxioCam MRc, Zeiss) and imaged with Zen pro 2.0 imaging software (Zeiss).

### T cell stimulation

Whole brains and spinal cords were mashed in digestion buffer (10 mM HEPES, 150 mM NaCl, 1 mM MgCl2, 5 mM KCl, and 1.8 mM CaCl2 in HBSS buffer). To dissociate cells from their resident tissues, collagenase D (Roche) was added to a final concentration of 1 mg/mL and DNAsel (Roche) to a final concentration of 60 μg/mL to each sample. CNS samples were incubated at 37°C for 30 minutes, mixed with a pipette tip, and reincubated for an additional 15 minutes. Upon removal, a final concentration of 1 mM EDTA pH 8.0 was added to each sample and incubated at room temperature for 10 minutes. Samples were then filtered through a 70-μm filter and washed twice with ice-cold PBS. Cells were resuspended into a 30% Percoll (GE Healthcare) solution and centrifuged to separate the fat from the cells. Collected lymphocytes were washed twice in ice-cold PBS and resuspended in complete RPMI (10% FBS from Gibco, Thermo Fisher Scientific; l-glutamine from MilliporeSigma, sodium pyruvate from MilliporeSigma, penicillin from MilliporeSigma, streptomycin from MilliporeSigma, HEPES pH 7.0 from Gibco, Thermo Fisher Scientific; and β-mercaptoethanol from Gibco, Thermo Fisher Scientific, in RPMI-1640 medium from MilliporeSigma). Whole brains and spinal cords were dissected and digested as described earlier. Cells were counted with a hemocytometer and plated at a density of 250,000 cells/well. Following incubation at 37°C for 5 hours with an intracellular cytokine restimulation buffer (PMA [MilliporeSigma, stock 500 μg/mL] used at 1:100,000; ionomycin [MilliporeSigma, stock 0.5 mg/mL] used at 1:1000; and Brefeldin A [eBioscience, Thermo Fisher Scientific, stock 100x] used at 1:1000 in complete RPMI), cells were collected, washed twice in ice-cold PBS, and stained for flow cytometry.

### Flow cytometry

Single-cell suspensions from CNS tissues were stained for viability with aqua, washed, and subsequently surface-stained with a panel of fluorescently-conjugated antibodies against CD45.1 (A20), CD3 (17A2), CD4 (RM4-5), CD19 (1D3), B220 (RA3-6B2), Gr-1 (RB6-8C5) or Ly6G (1A8), Ly6C (HK1.4), CD11b (M1/70), and CD11c (N418). Cell suspensions from T cell stimulation were surfaced-stained for CD4 (RM4-5), permeabilized with CytoFix/CytoPerm (BD Biosciences) for 20 minutes at 4°C, and subsequently stained intracellularly with GM-CSF (eBioscience), IFNγ (eBioscience), and IL-17A (eBioscience) (**Supp. Table 1**). Cells were acquired on a BD LSR X20 using FACS DIVA software.

### Single-cell isolation from leptomeningeal and cortical dissections

Mice were sacrificed by CO2 asphyxiation were decapitated, and skin overlying the skull was removed. Skull caps were carefully separated from the brain to remove dura mater and brains were transferred to petri dishes containing 1mL of ice-cold PBS. Under a dissection microscope, leptomeninges were removed from the brainstem, cerebellum, ventricles, hypothalamus, and cortex into digestion buffer (10 mM HEPES, 150 mM NaCl, 1 mM MgCl2, 5 mM KCl, and 1.8 mM CaCl2 in HBSS buffer) containing DNAse I (60μg/mL; Roche) and collagenase D (1mg/mL; Roche). Cortices were removed by first bisecting the brain along the sagittal plane to expose the corpus callosum, followed by removal of the brainstem, midbrain, cerebellum, and hypothalamus. Finally, cortices were isolated by dissecting away the corpus callosum and thalamus, and placed into digestion buffer. Cortices were mechanically dissociated by finely chopping via scalpel blade and then subjected to digestion with DNAse I and collagenase D for 30 minutes at 37C, 5% CO_2_. Cells from both leptomeninges and cortical digestions were subjected to 30% Percoll (GE Healthcare) gradient purification before downstream applications such as scRNAseq.

### Single-cell RNA sequencing

Prior to euthanasia, old and young EAE and naïve SJL/J mice were injected intravenously with 3μg of anti-CD45-PE (eBioscience) to label blood-derived and blood vessel-adjacent immune cells. Leptomeninges and cortices were dissected and single-cell suspensions were prepared as previously described. Cells from each individual mouse were stained using BioLegend TotalSeq B hashtags (Hashtags 1-4) and an anti-PE oligonucleotide barcode (BioLegend). Cells from each compartment (*i.e*. leptomeninges) were mixed in a 1:1 ratio, resuspended to a concentration of 1,300 cells/μL, and submitted for sequencing on the 10X Genomics platform using 5’ chemistry at the Princess Margaret Genomics Centre in Toronto, ON, Canada.

### Single molecular array assay for neurofilament light chain

The amount of NfL in mouse serum was quantified with a single-molecule array (Simoa) NF-light assay (Quanterix, Billerica, USA). In brief, magnetic beads were conjugated with monoclonal capture antibodies (mAB47:3, UmanDiagnostics), incubated with diluted mouse serum (1:8 or 1:16 dilution) and biotinylated detection antibodies (mAB2:1, UmanDiagnostics). Upon adding streptavidin-conjugated β-galactosidase (Quanterix), Resorufin β-D-galactopyranoside (Quanterix) was added for detection. The experiment was performed on a Simoa HD-X Analyzer (Quanterix). The assay was performed in duplicates and the mean of the two measured sNfL values per sample is reported.

### Preparation of samples for MRI imaging

Mice destined for MRI imaging were sacrificed by CO2 asphyxiation and transcardially perfused using a peristaltic pump at a rate of 1mL/min. Mice were first perfused with 40mL of PBS containing 2mM ProHance (Bracco Diagnostics) and 400USP heparin (Fresnius Kabi), followed by 30mL of PBS containing 2mM ProHance and 4% paraformaldehyde (EMS). Skulls were decapitated and placed into PBS containing 2mM ProHance and 4% paraformaldehyde (EMS). After an overnight incubation at 4C, skulls were transferred to PBS containing 2mM ProHance with 0.02% sodium azide (Fisher). Following 30-day incubation, skulls were scanned for MRI at the Mouse Imaging Centre in The Centre for Phenogenomics in Toronto, Canada.

### Anatomical image acquisition

A multi-channel 7 Tesla MRI scanner (Agilent Inc., Palo Alto, CA) was used to image brains in skulls. Sixteen samples were imaged in parallel using a custom-built 16-coil solenoid array^53^. To acquire anatomical images, the following scan parameters were used: T2W 3D FSE cylindrical k-space acquisition sequence, TR/TE/ETL = 350 ms/12 ms/6, two averages, FOV/matrix-size = 20 × 20 × 25 mm/504 × 504 × 630, total-imaging-time = 14 h. The resulting anatomical images had an isotropic resolution of 40μm voxels^54^.

### MRI registration and analysis

To assess any changes to the mouse brains due to age and treatment, all anatomical brain images were registered together using the mni_autoreg^55^ and ANTS (advanced normalizations tools)^56^ toolkits. The resulting consensus average and jacobian determinants were used to quantifying volumetric differences between each MR image and the average. The MAGeT pipeline^57^ was used to segment images using a published classified MRI.

### Post-mortem tissue retrieval

Tissue blocks for this study were obtained from the Netherlands Brain Bank (NBB; Amsterdam, The Netherlands). For the characterization of leptomeningeal immune cells, sample from twenty-seven donors with progressive (primary progressive, PP, or secondary progressive, SP) MS tissues were selected based on the presence of leptomeninges adjacent to cortex in the tissue blocks. For the characterization of MS subcortical white matter lesions, all available archived formalin-fixed paraffin-embedded (FFPE) tissue blocks for each of 27 MS patients (range 5-72 blocks, median: 30 blocks per donor) were analyzed (see **Supp. Table 2**). Tissue blocks were dissected based on the identification of lesions as guided by macroscopical examination and/or by post-mortem MRI (since 2001) of 1cm-thick coronal brain slices^58^. The tissue blocks used for the analysis of leptomeningeal inflammation and subpial demyelination performed in this study were dissected from the supratentorial cortex at locations that included the occipital or the parietal or the temporal or the frontal lobes.

Detailed clinical-pathological and demographic data of all donors are provided in **Supp. Table 2**. The age at the time of death of MS patients ranged from 41 to 81 years (median: 58 years) with a mean post-mortem delay of 8 hours and 31 minutes (SD, ± 1 hour 42 minutes). The age at the time of death of the non-neurological controls ranged from 49 to 99 years (median: 63.5 years) with a mean post-mortem delay of 9 hours 18 minutes (SD, ± 8 hours 37 minutes). The clinical diagnosis of MS and its clinical course were determined by a certified neurologist and confirmed by a certified neuropathologist based on the neuropathological analysis of the patient’s brain autopsy.

### Neuropathological techniques and immunohistochemistry

For the classification of cortical grey matter lesions, sections were stained by immunohistochemistry for the proteolipid protein (PLP) marker of myelin. For the identification of neutrophils, sections were stained with hematoxylin-eosin (H&E). Leptomeningeal immune cells were identified by immunohistochemistry for CD3 to detect T cells and CD20 to detect B cells (**Supp. Table 1**).

Immunohistochemistry was performed as previously described^59,60^. Sections of 7μm thickness were cut from formalin-fixed paraffin-embedded tissue blocks, collected on Superfrost Plus glass slides (VWR international; Leuven, Belgium) and dried overnight at 37°C. Sections were deparaffinized in xylene (2 x 15 minutes) and rehydrated through a series (100%, 70%, 50%) of ethanol. Endogenous peroxidase activity was blocked by incubation in methanol (Merck KGaA, Darmstadt, Germany) with 0.3% H2O2 (Merck KGaA) for 20 minutes at room temperature (RT). Sections were then rinsed in PBS and pre-treated with microwave antigen retrieval (3 minutes at 900W followed by 10 minutes at 90W) in either 0.05M tris buffered saline (TBS, pH 7.6) or 10 mM Tris/1 mM ethylenediaminetetraacetic acid (EDTA) buffer pH 9.0 (**Supp. Table 1**).

Sections were incubated overnight at 4°C in the appropriate primary antibody (**Supp. Table 1**) diluted in Normal Antibody Diluent (Immunologic, Duiven, The Netherlands) and the next day with the BrightVision poly-HRP-Anti Ms/Rb/Rt IgG biotin-free (diluted 1:1 in PBS, ImmunoLogic) for 30 minutes at RT. The immunostaining was visualized with 3,3’-diaminobenzidinetetrahydrochloride (DAB, Vector Laboratories) for 4 minutes at RT and sections were counterstained with haematoxylin (Sigma Chemie GmbH, Steinheim, Germany), dehydrated in ethanol and mounted with Pertex (Histolab, Gothenburg, Sweden).

### Quantification of leptomeningeal inflammation

Leptomeningeal segments were randomly selected for imaging at 20x magnification with a light microscope (Olympus BX41TF, Zoeterwoude, the Netherlands) connected to the Cell D software (Olympus, Zoeterwoude, the Netherlands). Immune cells were quantified in leptomeningeal areas that were adjacent to type III (subpial) grey matter lesions (GML) and in areas that were adjacent to normal appearing grey matter (NAGM). A total of 71±20% (mean±SD) of intact leptomeninges were available for scoring in the MS cohort and 40±8% (mean±SD) of intact leptomeninges were available for scoring in the NNC cohort. CD20^+^ B cell counts were done over a total leptomeningeal area of 47.53 mm^2^ from MS patients, of which 37.02 mm^2^ was adjacent to GMLs and 10.51 mm^2^ was adjacent to NAGM; and 5.652 mm^2^ of leptomeningeal area was adjacent to non-neurological control cortex. CD3^+^ T cell counts were done in a total leptomeningeal area of 60.97 mm^2^ from patients, of which 48.3 mm^2^ was adjacent to GMLs and 12.67mm^2^ was adjacent to NAGM; and 6.807 mm^2^ of leptomeningeal area was adjacent to non-neurological control cortex. The leptomeningeal area (in mm^2^) was measured using the “measurement” function of the Image Pro Plus 7.0 imaging software (MediaCybernetics, Rockville, MD, USA). Cell numbers were expressed as mean number per mm of intact leptomeninges.

### Study approval

All post-mortem human tissue was collected with informed consent for the use of material and clinical data for research purposes, in compliance with ethical guidelines of the Vrij Universiteit and Netherlands Brain Bank, Amsterdam, The Netherlands (Reference 2009/148). In addition, the University of Toronto Research Ethics Board (REB) granted approval for conducting histology on all post-mortem human tissue (study number: 36850). All animal experiments were conducted in accordance with institutional guidelines, with ethical approval from the University of Toronto Faculty of Medicine Animal Care Committee.

### Data analysis and statistics

Unless otherwise stated, all statistical tests were run using GraphPad Prism v8.0. All quantification data subjected to Shapiro-Wilk normality test. Only *p*-values <0.05 were considered significant. Flow cytometry data was analysed using FlowJo v17.0. Sequencing RNA sequencing data was processed using the CellRanger (v6.1.1) feature barcoding pipeline and read matrices were analyzed in R (v3.0) using the Seurat package (v3.0)^30^.

## References

1. D’Amico, E. et al. Late-onset and young-onset relapsing-remitting multiple sclerosis: evidence from a retrospective long-term follow-up study. Eur. J. Neurol. 25, 1425–1431 (2018).

2. Engelhardt, B. et al. Vascular, glial, and lymphatic immune gateways of the central nervous system. Acta Neuropathol. 132, 317–338 (2016).

3. Scalfari, A., Neuhaus, A., Daumer, M., Ebers, G. C. & Muraro, P. A. Age and disability accumulation in multiple sclerosis. Neurology 77, 1246–1252 (2011).

4. Calabrese, M. et al. Exploring the origins of grey matter damage in multiple sclerosis. Nat. Rev. 16, 147–158 (2015).

5. Tutuncu, M. et al. Onset of progressive phase is an age-dependent clinical milestone in multiple sclerosis. Mult. Scler. 19, 188–198 (2013).

6. Freilich, J. et al. Characterization of annual disease progression of multiple sclerosis patients: A population-based study. Mult. Scler. 24, 786–794 (2018).

7. Confavreux, C. & Vukusic, S. Accumulation of irreversible disability in multiple sclerosis: from epidemiology to treatment. Clin. Neurol. Neurosurg. 108, 327–332 (2006).

8. Chard, D. & Miller, D. Grey matter pathology in clinically early multiple sclerosis: evidence from magnetic resonance imaging. J. Neurol. Sci. 282, 5–11 (2009).

9. Lucchinetti, C. F. et al. Inflammatory cortical demyelination in early multiple sclerosis. N. Engl. J. Med. 365, 2188–2197 (2011).

10. Kutzelnigg, A. et al. Cortical demyelination and diffuse white matter injury in multiple sclerosis. Brain 128, 2705–2712 (2005).

11. Fisniku, L. K. et al. Gray matter atrophy is related to long-term disability in multiple sclerosis. Ann. Neurol. 64, 247–254 (2008).

12. Haider, L. et al. Cortical involvement determines impairment 30 years after a clinically isolated syndrome. Brain 144, 1384–1395 (2021).

13. Scalfari, A. et al. The cortical damage, early relapses, and onset of the progressive phase in multiple sclerosis. Neurology 90, e2099–e2106 (2018).

14. Lünemann, J. D., Ruck, T., Muraro, P. A., Bar’Or, A. & Wiendl, H. Immune reconstitution therapies: concepts for durable remission in multiple sclerosis. Nat. Rev. Neurol. 16, 56–62 (2020).

15. Stys, P. K., Zamponi, G. W., Van Minnen, J. & Geurts, J. J. G. Will the real multiple sclerosis please stand up? Nat. Rev. Neurosci. 13, 507–514 (2012).

16. Serafini, B., Rosicarelli, B., Magliozzi, R., Stigliano, E. & Aloisi, F. Detection of ectopic B-cell follicles with germinal centers in the meninges of patients with secondary progressive multiple sclerosis. Brain Pathol. 14, 164–174 (2004).

17. Howell, O. W. et al. Extensive grey matter pathology in the cerebellum in multiple sclerosis is linked to inflammation in the subarachnoid space. Neuropathol. Appl. Neurobiol. 41, 798–813 (2015).

18. Absinta, M. et al. Gadolinium-based MRI characterization of leptomeningeal inflammation in multiple sclerosis. Neurology 85, 18–28 (2015).

19. Howell, O. W. et al. Meningeal inflammation is widespread and linked to cortical pathology in multiple sclerosis. Brain 134, 2755–2771 (2011).

20. Magliozzi, R. et al. Meningeal B-cell follicles in secondary progressive multiple sclerosis associate with early onset of disease and severe cortical pathology. Brain 130, 1089–1104 (2007).

21. Magliozzi, R. et al. A Gradient of neuronal loss and meningeal inflammation in multiple sclerosis. Ann. Neurol. 68, 477–493 (2010).

22. Magliozzi, R. et al. Inflammatory intrathecal profiles and cortical damage in multiple sclerosis. Ann. Neurol. 83, 739–755 (2018).

23. Pikor, N. B. et al. Integration of Th17- and Lymphotoxin-Derived Signals Initiates Meningeal-Resident Stromal Cell Remodeling to Propagate Neuroinflammation. Immun. (Cambridge, Mass.) 43, 1160–1173 (2015).

24. Ward, L. A. et al. Siponimod therapy implicates Th17 cells in a preclinical model of subpial cortical injury. JCI insight 5, (2020).

25. Calabrese, M. et al. The changing clinical course of multiple sclerosis: a matter of gray matter. Ann. Neurol. 74, 76–83 (2013).

26. Magliozzi, R. et al. A Gradient of neuronal loss and meningeal inflammation in multiple sclerosis. Ann. Neurol. 68, 477–493 (2010).

27. Kapoor, R. et al. Serum neurofilament light as a biomarker in progressive multiple sclerosis. Neurology 95, 436–444 (2020).

28. Kiljan, S. et al. Cortical axonal loss is associated with both gray matter demyelination and white matter tract pathology in progressive multiple sclerosis: Evidence from a combined MRI-histopathology study. Mult. Scler. 27, 380–390 (2021).

29. Geurts, J. J. G., Calabrese, M., Fisher, E. & Rudick, R. A. Measurement and clinical effect of grey matter pathology in multiple sclerosis. Lancet Neurol. 11, 1082–1092 (2012).

30. Stuart, T. et al. Comprehensive Integration of Single-Cell Data. Cell 177, 1888–1902.e21 (2019).

31. Wu, F., Cao, W., Yang, Y. & Liu, A. Extensive infiltration of neutrophils in the acute phase of experimental autoimmune encephalomyelitis in C57BL/6 mice. Histochem. Cell Biol. 133, 313–322 (2010).

32. Ahmed, S. M. et al. Accumulation of meningeal lymphocytes, but not myeloid cells, correlates with subpial cortical demyelination and white matter lesion activity in progressive MS patients. medRxiv 2021.12.20.21268104 (2021) doi:10.1101/2021.12.20.21268104.

33. Kipp, M., Nyamoya, S., Hochstrasser, T. & Amor, S. Multiple sclerosis animal models: a clinical and histopathological perspective. Brain Pathol. 27, 123–137 (2017).

34. Peferoen, L. A. N. et al. Ageing and recurrent episodes of neuroinflammation promote progressive experimental autoimmune encephalomyelitis in Biozzi ABH mice. Immunology 149, 146–156 (2016).

35. Absinta, M., Lassmann, H. & Trapp, B. D. Mechanisms underlying progression in multiple sclerosis. Curr. Opin. Neurol. 33, 277–285 (2020).

36. Ciotti, J. R. & Cross, A. H. Disease-Modifying Treatment in Progressive Multiple Sclerosis. Curr. Treat. Options Neurol. 20, (2018).

37. Li, J. L. Y. et al. Neutrophils Self-Regulate Immune Complex-Mediated Cutaneous Inflammation through CXCL2. J. Invest. Dermatol. 136, 416–424 (2016).

38. Capucetti, A., Albano, F. & Bonecchi, R. Multiple Roles for Chemokines in Neutrophil Biology. Front. Immunol. 11, 1259 (2020).

39. Werneburg, S. et al. Targeted Complement Inhibition at Synapses Prevents Microglial Synaptic Engulfment and Synapse Loss in Demyelinating Disease. Immunity 52, 167–182.e7 (2020).

40. Ramaglia, V. et al. Complement-associated loss of CA2 inhibitory synapses in the demyelinated hippocampus impairs memory. Acta Neuropathol. 142, 643 (2021).

41. De Bondt, M., Hellings, N., Opdenakker, G. & Struyf, S. Neutrophils: Underestimated Players in the Pathogenesis of Multiple Sclerosis (MS). Int. J. Mol. Sci. 21, 1–25 (2020).

42. Yamasaki, R. et al. Differential roles of microglia and monocytes in the inflamed central nervous system. J. Exp. Med. 211, 1533–1549 (2014).

43. Lapinet, J. A., Scapini, P., Calzetti, F., Pérez, O. & Cassatella, M. A. Gene expression and production of tumor necrosis factor alpha, interleukin-1beta (IL-1beta), IL-8, macrophage inflammatory protein 1alpha (MIP-1alpha), MIP-1beta, and gamma interferon-inducible protein 10 by human neutrophils stimulated with group B meningococcal outer membrane vesicles. Infect. Immun. 68, 6917–6923 (2000).

44. Thom, S. R. et al. Neutrophil microparticle production and inflammasome activation by hyperglycemia due to cytoskeletal instability. J. Biol. Chem. 292, 18312 (2017).

45. Gijbels, K. et al. Gelatinase B is present in the cerebrospinal fluid during experimental autoimmune encephalomyelitis and cleaves myelin basic protein. J. Neurosci. Res. 36, 432–440 (1993).

46. Yu, G., Zheng, S. & Zhang, H. Inhibition of myeloperoxidase by N-acetyl lysyltyrosylcysteine amide reduces experimental autoimmune encephalomyelitis-induced injury and promotes oligodendrocyte regeneration and neurogenesis in a murine model of progressive multiple sclerosis. Neuroreport 29, 208–213 (2018).

47. Norden, D. M. & Godbout, J. P. Review: Microglia of the aged brain: Primed to be activated and resistant to regulation. Neuropathology and Applied Neurobiology vol. 39 19–34 (2013).

48. Li, K., Li, J., Zheng, J. & Qin, S. Reactive astrocytes in neurodegenerative diseases. Aging and Disease vol. 10 664–675 (2019).

49. Hickman, S., Izzy, S., Sen, P., Morsett, L. & El Khoury, J. Microglia in neurodegeneration. Nature Neuroscience vol. 21 (2018).

50. Torre, S. et al. USP15 regulates type I interferon response and is required for pathogenesis of neuroinflammation. Nat. Immunol. 18, 54–63 (2017).

51. Giuliani, F. et al. Additive effect of the combination of glatiramer acetate and minocycline in a model of MS. J. Neuroimmunol. 158, 213–221 (2005).

52. Galicia, G. et al. Isotype-Switched Autoantibodies Are Necessary To Facilitate Central Nervous System Autoimmune Disease in Aicda -/- and Ung -/-Mice. J. Immunol. 201, 1119–1130 (2018).

53. Bock, N. A., Nieman, B. J., Bishop, J. B. & Henkelman, R. M. In vivo multiple-mouse MRI at 7 Tesla. Magn. Reson. Med. 54, 1311–1316 (2005).

54. Spencer Noakes, T. L., Henkelman, R. M. & Nieman, B. J. Partitioning k-space for cylindrical three-dimensional rapid acquisition with relaxation enhancement imaging in the mouse brain. NMR Biomed. 30, (2017).

55. Collins, D. L., Neelin, P., Peters, T. M. & Evans, A. C. Automatic 3D Intersubject Registration of MR Volumetric Data in Standardized Talairach Space. J. Comput. Assist. Tomogr. 18, 192–205 (1994).

56. Avants, B. B. et al. A reproducible evaluation of ANTs similarity metric performance in brain image registration. Neuroimage 54, 2033–2044 (2011).

57. Chakravarty, M. M. et al. Performing label-fusion-based segmentation using multiple automatically generated templates. Hum. Brain Mapp. 34, 2635–2654 (2013).

58. Luchetti, S. et al. Progressive multiple sclerosis patients show substantial lesion activity that correlates with clinical disease severity and sex: a retrospective autopsy cohort analysis. Acta Neuropathol. 135, 511–528 (2018).

59. Frischer, J. M. et al. The relation between inflammation and neurodegeneration in multiple sclerosis brains. Brain 132, 1175 (2009).

60. Michailidou, I. et al. Complement C1q-C3-associated synaptic changes in multiple sclerosis hippocampus. Ann. Neurol. 77, 1007–1026 (2015).

