## Supplemental material for "Age-related susceptibility to grey matter demyelination and neurodegeneration is associated with meningeal neutrophil accumulation in an animal model of Multiple Sclerosis"

### Supplemental Figures

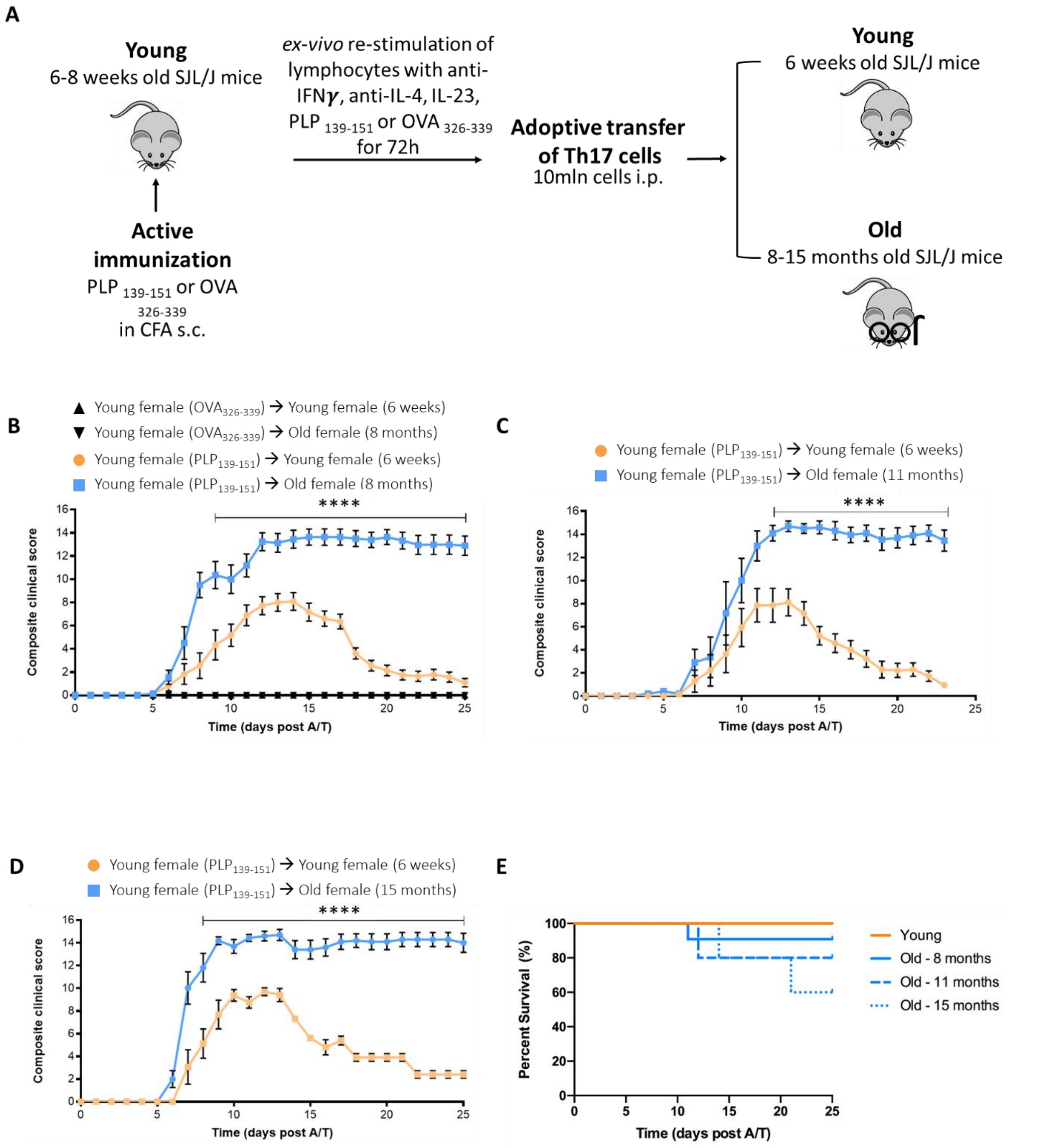


**Supplemental figure 1. The non-remitting phenotype of SJL/J A/T EAE is reproducible. (A)** Induction protocol for A/T EAE in young (6 week) *vs* old (8-15 month) SJL/J mice. **(B)** Clinical course of A/T EAE using cells from $PLP_{139-151}$- or $OVA_{326-339}$-primed donors. Only mice receiving cells from $PLP_{139-151}$-primed donors developed clinical disease (young n=18, old n=11), while mice receiving cells from $OVA_{326-339}$-primed donors remained asymptomatic (young n=4, old n=3). **(C)** Clinical course of A/T EAE in 6 week (n=8) vs 11 month-old (n=8) recipients. **(D)** Clinical course of A/T EAE in 6 week (n=8) vs 15 month-old (n=8) recipients. **(E)** Kaplan-Meier curve showing age-dependent survival in SJL/J A/T EAE mice. Stats for B, C, D = Two-way ANOVA with Bonferroni correction for multiple comparisons. Experiments were all performed at U of T. Error bars indicate mean with SEM.

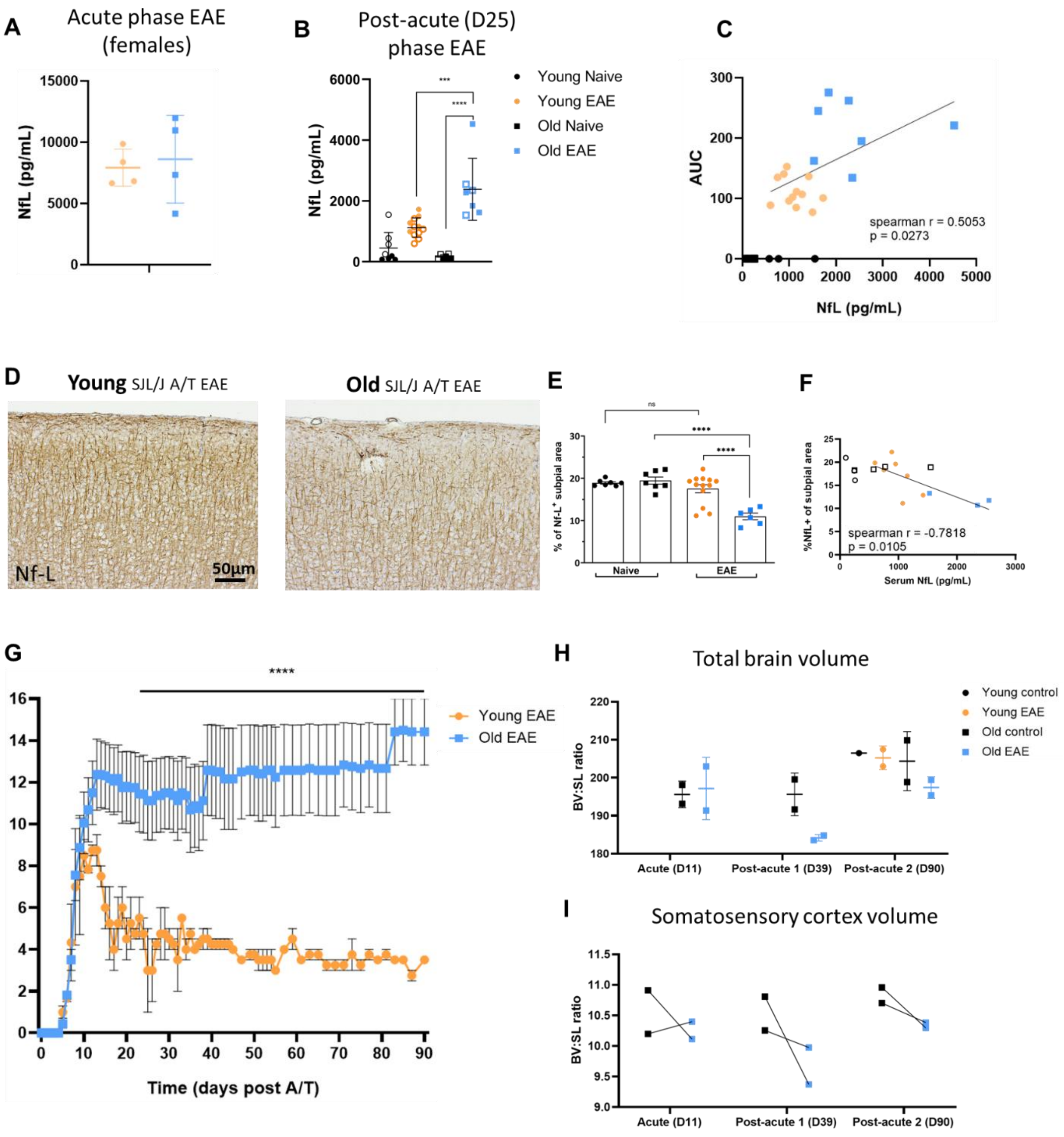

A
Acute phase EAE
(females)
NfL (pg/mL)
B
Post-acute (D25)
phase EAE
Young Naive
Young EAE
Old Naive
Old EAE
C
AUC
spearman r = 0.5053
p = 0.0273
NfL (pg/mL)
D
Young SJL/J A/T EAE
Old SJL/J A/T EAE
Nf-L
50µm
E
ns
Naive
EAE
F
%NfL+ of subpial area
spearman r = -0.7818
p = 0.0105
Serum NfL (pg/mL)
G
Young EAE
Old EAE
Time (days post A/T)
H
Total brain volume
Young control
Young EAE
Old control
Old EAE
BV:SL ratio
Acute (D11)
Post-acute 1 (D39)
Post-acute 2 (D90)
I
Somatosensory cortex volume
BV:SL ratio
Acute (D11)
Post-acute 1 (D39)
Post-acute 2 (D90)

**Supplemental Figure 2. Ageing induces accumulation of NfL in SJL/J A/T EAE mice and loss of brain volume at post-acute disease stage.** Sera of old vs young naïve and SJL/J A/T EAE mice were assayed for neurofilament light chain (sNfL) on the Quanterix single molecule array (Simoa) platform. **(A)** At the acute (D11) phase of disease, no difference between the groups was noted. **(B)** At post-acute (D25) disease stage, a difference in sNfL levels was noted between both females (filled points) and males (open points). Stats = Kruskal-Wallis with correction for multiple comparisons. **(C)** Correlation analysis of clinical severity represented by area under the curve (AUC) with levels of serum NfL revealed a significant positive correlation. Stats by Spearman's correlation test. **(D)** Representative immunohistochemistry staining for NfL in the cortex of old vs young SJL/J A/T EAE mice. **(E)** Quantification of IHC staining for NfL using %NfL$^+$ area in the subpial cortex. Stats by Kruskal-Wallis with correction for multiple comparisons revealed no difference between groups. **(F)** Correlation analysis of IHC quantification (represented by %NfL$^+$ area) with levels of sNfL revealed a significant negative correlation. Stats by Spearman's correlation test. **(G)** Clinical course of old vs young SJL/J A/T EAE mice followed for 90 days post-adoptive transfer. Stats by two-way ANOVA with Bonferroni's correction for multiple comparisons. Old mice were sacrificed at acute (D11) and two post-acute stages (D39, D90). Young mice were sacrificed at the last post-acute disease stage (D90). Skulls were sent for MRI scanning. **(H)** Brain volume of each mouse measured by MRI normalized to skull length. **(I)** Cumulative volumes of somatosensory cortex regions of each old mouse normalized by skull length. Error bars indicate mean with SD unless otherwise stated.

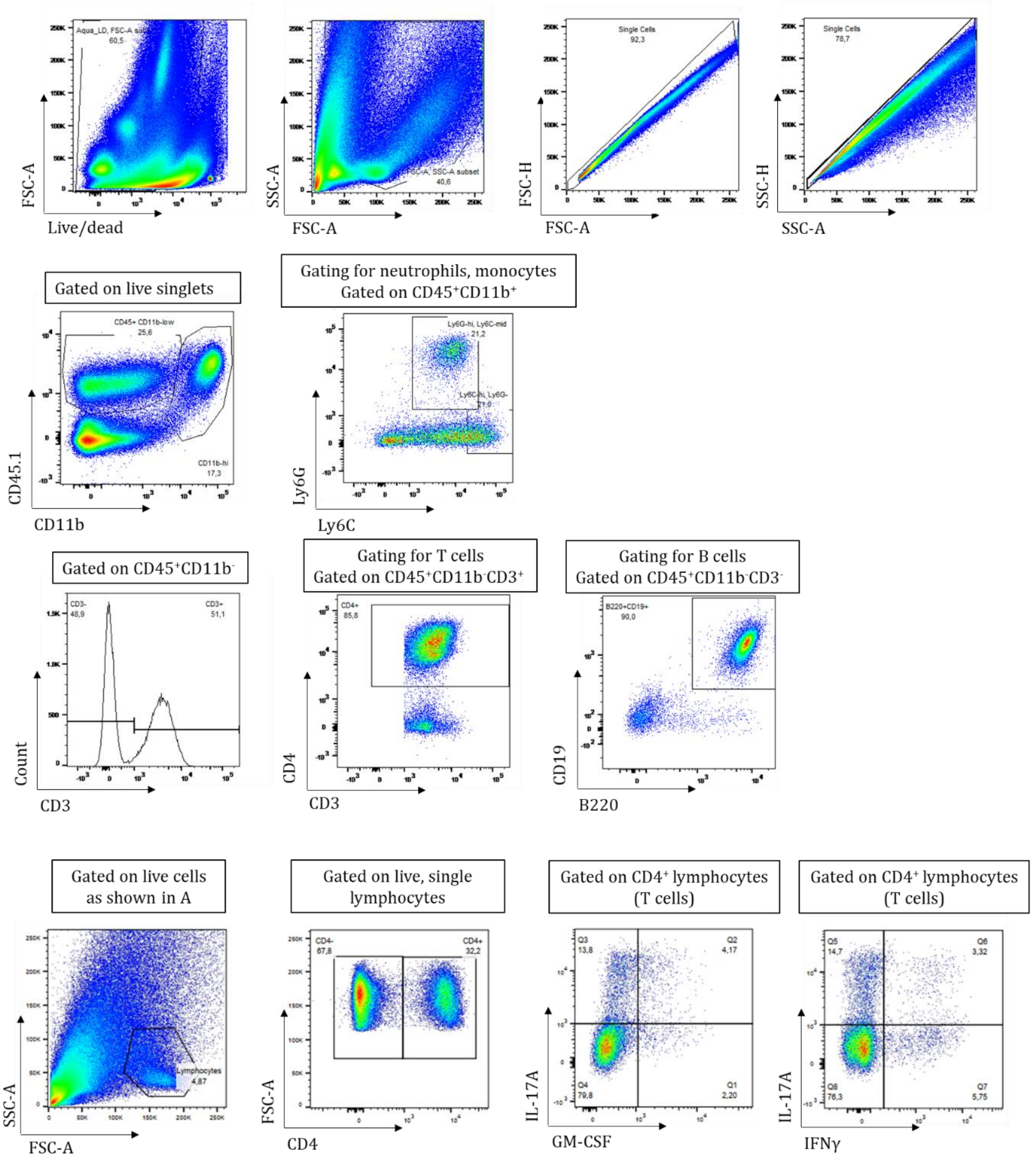


**Supplemental Figure 3. Gating strategies for identification of T cells, B cells, neutrophils, and cytokine-producing cells.** **(A)** Pre-gating for live, single cells. **(B)** Gating for identification of neutrophils (CD45+CD11b+Ly6G+Ly6C-mid). **(C)** Gating for identification of T cells (CD45+CD11b-CD3+CD4+) and B cells (CD45+CD11b-CD3-B220+CD19+). **(D)** Pre-gating for lymphocytes, following by identification of CD4+ cytokine producing cells.

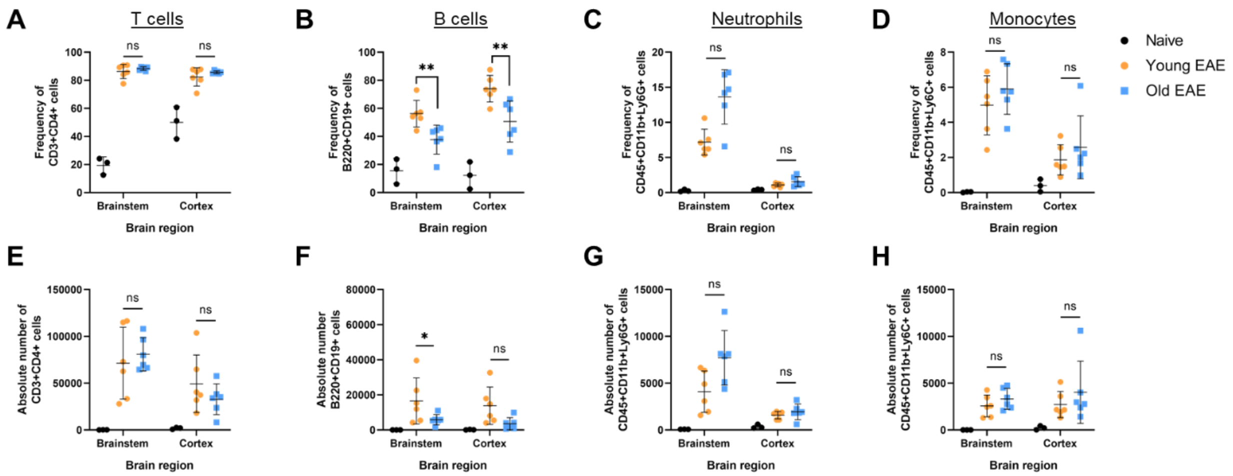


**Supplemental Figure 4. Flow cytometry of cortex and brainstem from old vs young SJL/J A/T EAE mice reveals differences in B cell composition only.** Brainstem and cortex was separately dissected from old vs young SJL/J A/T EAE mice at the acute timepoint and subjected to analysis by flow cytometry. Despite no change in **(A, E)** T cell, **(C, G)** neutrophils, or **(D,H)** monocytes in number or frequency, we observed a decrease in density of **(B, F)** B cells. Stats by one-way ANOVA (absolute number) or Kruskal-Wallis (frequency) with correction for multiple comparisons. Error bars indicate mean with SD.

### Supplemental Tables

**Supplemental table 1. Antibodies used for immunohistochemistry, immunofluorescence, and flow cytometry.**

| Primary Antibody | *Clone* | Target | Dilution | Antigen Retrieval | Source |
|---|---|---|---|---|---|
| *Immunohistochemistry* | | | | | |
| PLP | mAb, plpc1 | Proteolipid protein (Myelin) | 1:100 | Tris-EDTA | Bio-Rad, MCA839G |
| Iba-1 | mAb, EPR16589 | Ionized alcium binding adaptor molecule 1 (microglia/macrophages) | 1:4000 | Tris-EDTA | Abcam, ab92305 |
| GFAP | pAb | Glial fibrillary acidic protein (astrocytes) | 1:4000 | Tris-EDTA | Dako, Z0334 |
| Synaptophysin | mAb, SY38 | Synapses | 1:2500 | Tris-EDTA | ThermoFisher, MA1-213 |
| Pan-Neurofilament | pAb | Axons | 1:100 | Tris-EDTA | Abcam, ab204893 |
| Neurofilament light chain | mAb, 8A1 | Axons | 1:100 | Tris-EDTA | Santa Cruz Biotech, sc-20012 |
| CD3 | mAb, SP7 | T cells | 1:100 | Citrate | Abcam, ab16669 |
| CD20 | mAb, L26 | B cells | 1:100 | Citrate | Abcam, ab9475 |
| *Immunofluorescence* | | | | | |
| CD45-AF594* | mAb, 30-F11 | Lymphocytes and myeloid cells | 1:200 | - | Biolegend, 103144 |
| CD3-AF594* | mAb, 17A2 | T cells | 1:100 | - | Biolegend, 100240 |
| B220-AF488* | mAb, RA3-6B2 | B cells | 1:100 | - | eBioscience 53-0452-82 |
| Ly6G-PE* | mAb, 1A8 | Neutrophils | 1:100 | - | eBioscience, 12-9668-82 |
| Fibronectin* | mAb, GW20021F | Extracellular matrix | 1:100 | - | Sigma, 53-0452-83 |
| *Flow cytometry* | | | | | |
| CD45.1-BUV395 | mAb, A20 | Lymphocytes and myeloid cells | 1:100 | - | BD Horizon, 565212 |
| CD19-eF450 | mAb, 1D3 | B cells | 1:200 | - | eBioscience, 48-0193-82 |
| B220-BV605 | mAb, RA3-6B2 | B cells | 1:200 | - | Biolegend, 103244 |
| CD3-BV711 | mAb, 17A2 | T cells | 1:200 | - | Biolegend, 100241 |
| Ly6C-PerCP-Cy5.5 | mAb, HK1.4 | Monocytes | 1:200 | - | eBioscience, 45-5932-82 |

| | | | | | |
|---|---|---|---|---|---|
| Ly6G-PE | mAb, 1A8 | Neutrophils | 1:200 | - | eBioscience, 12-9668-82 |
| CD11b-PE-Cy7 | mAb, M1/70 | Myeloid cells | 1:200 | - | eBioscience, 25-0112-82 |
| CD4-APC | mAb, RM4-5 | MHC class II restricted T cells | 1:200 | - | eBioscience, 17-0042-82 |
| CD8α-PE-Cy7 | mAb, 53-6.7 | MHC class I restricted T cells | 1:200 | - | eBioscience, 25-0081-82 |
| CD11c-APC-Cy7 | mAb, N418 | Myeloid cells | 1:200 | - | Biolegend, 117324 |
| GM-CSF-FITC | mAb, MP1-22E9 | Granulocyte-macrophage colony-stimulating factor | 1:100 | - | eBioscience, 11-7331 |
| IL17a-PerCP-Cy5.5 | mAb, 17B7 | Interleukin 17 alpha | 1:100 | - | eBioscience, 45-7177 |
| IFNγ-PE-Cy7 | mAb, XMG1.2 | Interferon gamma | 1:200 | - | eBioscience, 25-7311 |
| Brefeldin A | | | 1:1000 | - | eBioscience, 00-4506-51 |
| Aqua fluorescein viability dye | | | 1:1000 | - | ThermoFisher, L34965 |

mAb, monoclonal antibody; pAb, polyclonal antibody; Tris-EDTA, 10mM Tris 1mM EDTA buffer pH 9.0; Citrate, 10mM citrate buffer pH 6.0; the asterisk indicates antibodies used on frozen sections

**Supplementary Table 2. Donor demographics.**

| Case | Sex | Age range (years) | PMD (h:min) | Type of MS | DD (years) | No. tissue blocks analysed for lesion characterization | COD |
|---|---|---|---|---|---|---|---|
| ***MS*** | | | | | | | |
| 1 | F | 48-81 | 08:40 | SPMS | 26 | 41 | Respiratory insufficiency to (uro)sepsis |
| 2 | F | 48-81 | 10:40 | SPMS | 29 | 30 | Euthanasia |
| 3 | F | 48-81 | 07:30 | SPMS | 34 | 27 | Euthanasia |
| 4 | F | 48-81 | 11:50 | PPMS | 22 | 32 | Respiratory failure with end stage MS |
| 5 | M | 48-81 | 08:15 | SPMS | 30 | 23 | Pneumonia, cachexia and dehydration |
| 6 | F | 48-81 | 09:05 | SPMS | 18 | 23 | Euthanasia |
| 7 | F | 48-81 | 08:35 | PPMS | 29 | 34 | Aspiration pneumonia |
| 8 | F | 48-81 | 05:45 | PPMS | 22 | 15 | Sepsis |
| 9 | F | 48-81 | 08:25 | SPMS | 11 | 8 | Natural death |
| 10 | M | 48-81 | 11:00 | SPMS | >12 | 38 | Exact cause unknown, infection 2 days prior to death |
| 11 | F | 48-81 | 08:40 | PPMS | 29 | 31 | Euthanasia |
| 12 | F | 48-81 | 08:25 | SPMS | 34 | 27 | Respiratory insufficiency secondary to pneumonia |
| 13 | M | 48-81 | 07:30 | - | - | 28 | - |
| 14 | F | 48-81 | 06:45 | PPMS | 29 | 5 | Cardiac asthma |
| 15 | F | 48-81 | 07:30 | SPMS | 39 | 23 | Bronchitis/ aspiration pneumonia |
| 16 | M | 48-81 | 10:45 | RRMS | 22 | 34 | Euthanasia |
| 17 | F | 48-81 | 08:00 | SPMS | 42 | 32 | Pneumonia |
| 18 | M | 48-81 | 06:20 | PPMS | 32 | 33 | Respiratory insufficiency |
| 19 | F | 48-81 | 10:05 | SPMS | 26 | 18 | Cardiovascular event and dehydration |
| 20 | F | 48-81 | 07:50 | SPMS | 50 | 72 | Euthanasia |
| 21 | M | 48-81 | 10:45 | RRMS | 22 | 34 | Euthanasia |
| 22 | F | 48-81 | 07:05 | SPMS | 34 | 26 | Cachexia with slowly progressive MS and metastatic breast cancer |
| 23 | F | 48-81 | 09:35 | PPMS | 22 | 25 | Cardiac asthma |
| 24 | F | 48-81 | 04:35 | SPMS | 34 | 30 | Aspiration pneumonia |
| 25 | M | 48-81 | 09:15 | - | - | 30 | - |
| 26 | M | 48-81 | 07:30 | PPMS | 32 | 30 | Respiratory failure due to pneumonia |
| 27 | F | 48-81 | 09:45 | SPMS | 35 | 34 | Euthanasia |
| ***Non-neurological Controls*** | | | | | | | |
| 28 | M | 49-99 | 06:15 | - | - | - | Euthanasia with Hopkin's lymphoma |
| 29 | F | 49-99 | 04:15 | - | - | - | - |
| 30 | M | 49-99 | 26:45 | - | - | - | Pulmonary infection |
| 31 | F | 49-99 | 07:30 | - | - | - | Infection e.c.i |
| 32 | M | 49-99 | 05:00 | - | - | - | - |
| 33 | F | 49-99 | 06:00 | - | - | - | Cachexia |
| 34 | M | 49-99 | 14:00 | - | - | - | Heart failure |
| 35 | F | 49-99 | 7:40 | - | - | - | - |
| 36 | M | 49-99 | 7:44 | - | - | - | - |

MS,multiple sclerosis; F, Female; M, Male; PMD, post-mortem delay; DD, disease duration; h:min, hours:minutes; COD, cause of death.
